# Molecular Genomic Insights into Melanoma Associated Proteins PRAME and BAP1

**DOI:** 10.1101/2024.03.05.583532

**Authors:** Debaleena Nawn, Sk. Sarif Hassan, Altijana Hromić-Jahjefendić, Tanishta Bhattacharya, Pallab Basu, Elrashdy M. Redwan, Debmalya Barh, Bruno Silva Andrade, Alaa A. Aljabali, Ángel Serrano-Aroca, Kenneth Lundstrom, Murtaza M. Tambuwala, Vladimir N. Uversky

## Abstract

**Background:** Melanoma, a worldwide widespread skin cancer with over 325,000 yearly incidences, demands a thorough understanding of its molecular components to create effective therapeutics. This study looks at the PRAME (cutaneous melanoma-associated antigen) and BAP1 (gene controlling gene-environment interactions) proteins, which are important in melanoma development and are important for understanding the molecular landscape of melanoma.

**Introduction:** While playing a crucial role in melanoma, the structural and functional characteristics of PRAME and BAP1 remain unidentified. This work tries to unravel their complexities by investigating conserved residues, sequence invariance, and other molecular characteristics that contribute to their importance in melanoma. Promising therapeutic targets for melanoma therapy are identified by analyzing these proteins at the molecular level.

**Methods:** The study makes extensive use of bioinformatics methods to analyze PRAME and BAP1, including sequence conservation, inherent disorder, polyglutamic acid presence, and polarity alterations. Established approaches are used to investigate residue changes and their effects on protein folding, aggregation, and interactions.

**Results:** PRAME and BAP1 conserved residues highlight their critical roles in protein function and interaction. Sequence invariance indicates the possibility of functional relevance and evolutionary conservation. In intrinsically disordered proteins (IDPRs), PRAME has enhanced intrinsic disorder and flexibility, whereas BAP1 has changed disorder-promoting residue sequences. Polyglutamic acid strings are found in both proteins, emphasizing their modulatory involvement in protein interactions. Protein folding and aggregation are influenced by polarity shifts, with a balanced distribution of acidic and basic residues preserving native structures. The ratios and distributions of amino acids, particularly neutral residues, have a profound influence on interactions and gene dysregulation.

**Conclusion:** PRAME and BAP1 structural and functional understanding pave the way for diagnostic and tailored treatment options in melanoma. Differences in residue alterations, polarity distributions, and amino acid ratios provide intriguing drug design options. This research contributes to a better knowledge of melanoma-associated two proteins, opening the path for novel diagnostic and therapy techniques in skin cancer and beyond.

## 1. Introduction

Melanoma or malignant melanoma is a type of cancer that originates in pigment-containing cells known as melanocytes. Melanoma remains one of the most fatal cancers since the turn of the century. When detected early, most patients are treated through local surgery after a sentinel lymph node biopsy [1]. The prevalence varies according to country, skin phenotype, and sun exposure. It usually affects youngsters and middle-aged female populations (those under the age of 50), although males beyond the age of 55 are more affected [1]. Melanoma is three times more prevalent in males compared to women by the age of 75. It has been reported that UV radiation is recognized to be the main trigger of malignant melanoma. The history of sunburn in childhood has been linked directly to the occurrence of melanoma. The number of melanocytic nevi, family history, and genetic background are all risk factors. It has also been reported that people who have previous cases of melanoma are prone to acquire numerous primary melanomas [1]. Other environmental variables, such as alcohol or tobacco use, have not been reported as a causative agent in melanoma development [2]. Various factors like the origin of the exposure to sunlight, the amount of cumulative UV exposure, the age at the time of diagnosis, the types of oncogenic drivers, and the mutational load have all been used in skin melanoma classification [2]. The major genetic causes are known to include B-Raf proto-oncogene (BRAF), neurofibromin 1 (NF1), and NRAS mutations, as well as a high mutational burden caused by UV radiation [3]. Periodic UV exposure is frequently linked to the BRAFV600E mutation and a reduced mutational burden. It is crucial to note that each melanoma subtype can develop from a variety of antecedent lesions, which can include a variety of gene alterations as well as distinct transformative phases [2]. Current medical treatments employ a variety of techniques. Most individuals with newly identified melanoma have early-stage illness and may be treated with surgical excision, which is usually curative. In addition to traditional surgical excision, certain therapy approaches include lymph node biopsies. Sadly, 10% of all melanoma cases are discovered at an advanced/late stage and have already spread to other parts of the body, including visceral and cerebral metastases [2, 3, 4, 5]. Recently it has been reported that several protein markers play a crucial role in diagnosis of melanoma and they are PRAME and BAP1. Preferentially expressed Antigen in Melanoma (PRAME) is a tumor-associated peptide discovered by studying the specificity of tumor-reactive T-cell clones obtained from a patient with metastatic cutaneous melanoma [6]. PRAME was later discovered to be expressed not only in cutaneous melanoma, but also in ocular melanoma and a variety of nonmelanocytic malignant neoplasms such as non-small cell lung cancer, breast carcinoma, renal cell carcinoma, ovarian carcinoma, leukemia, synovial sarcoma, cervical cancer, and myxoid liposarcoma [7, 8, 9]. Except for the testis, ovary, placenta, adrenals, and endometrium, no normal healthy tissues have been found to express PRAME [10]. The BRCA1 (BReast CAncer gene 1)-associated protein (BAP1) gene, on the other hand, produces a tumor suppressor protein that acts through numerous unidentified routes. BAP1 is found on chromosome 3p21.1, which is commonly deleted or mutated in various types of melanomas [11, 12]. BAP1 mutations include missense and nonsense changes, frame-shift deletions, non-frame-shift deletions, and four splice site changes. These results and connections are essential for better understanding of melanoma pathogenesis; nevertheless, further research is needed to understand how these genetic variants affect the diverse activities of this critical protein. It is unknown how these mutations affect BAP1’s functional proteomics and interactions with additional proteins [13]. The goal of this study is to provide information on the intrinsic regulatory regions that regulate these critical genes in melanoma. This study aims to expand the knowledge of melanoma biology and find fresh routes for targeted therapies by thoroughly investigating their functions and interactions.

## 2. Data acquisition

The protein sequences for Melanoma antigen preferentially expressed in tumors (PRAME) and Ubiquitin carboxyl-terminal hydrolase BAP1 (BAP1) were obtained from UniProt by conducting BLAST (by customizing the similarity percentage from 95% to 100%) searches using the human PRAME (UniProt ID: P78395) and human BAP1 (UniProt ID: Q92560) sequences as queries. It is important to note that none of the nine BAP1 proteins from various organisms were identical. Among the 25 PRAME sequences, 21 were found to be distinct and unique. The following sets of PRAME sequences were identical.

▪ {(tr|A0A096N6T6|PRAME_PAPAN), (tr|A0A2K5MV40|PRAME_CERAT), (tr|A0A8D2JTV4|PRAME_THEGE)}

▪ {(tr|A0A2K5TZS8|PRAME_MACFA) and (tr|F6QQN0|PRAME_MACMU)}

▪ {(tr|A0A2K6LQQ8|PRAME_RHIBE) and (tr|A0A2K6QD00|PRAME_RHIRO)}

The list of unique PRAME/BAP1 are presented in Table 1.

**Table 1:** List of 21 PRAME and 9 BAP1 proteins from various organisms with their associated Uniprot ID (Hyperlinked with respective Uniprot link)

| Serial No | UNIPOINT ID | Renamed as | Serial No | UNIPOINT ID | Renamed as | Serial No | UNIPOINT ID | Renamed as |
| --- | --- | --- | --- | --- | --- | --- | --- | --- |
| 1 | tr\|A0A2K6AQC0\|PRAME_MACNE | PRAME_1 | 12 | tr\|A0A2K6AQA6\|PRAME_MACNE | PRAME_12 | 1 | sp\|Q52L14\|BAP1_XENLA | BAP1_1 |
| 2 | tr\|A0A1D5R115\|PRAME_MACMU | PRAME_2 | 13 | tr\|A0A2K5TZS8\|PRAME_MACFA | PRAME_13 | 2 | sp\|Q66JB6\|BAP1_XENTR | BAP1_2 |
| 3 | tr\|A0A2K5MV09\|PRAME_CERAT | PRAME_3 | 14 | tr\|A0A096N6T6\|PRAME_PAPAN | PRAME_14 | 3 | sp\|Q99PU7\|BAP1_MOUSE | BAP1_3 |
| 4 | tr\|A0A2K5Y2X5\|PRAME_MANLE | PRAME_4 | 15 | tr\|A0A2K5Y2Y9\|PRAME_MANLE | PRAME_15 | 4 | sp\|A2VDM8\|BAP1_BOVIN | BAP1_4 |
| 5 | tr\|A0A2K6LQR2\|PRAME_RHIBE | PRAME_5 | 16 | tr\|H2QLB6\|PRAME_PANTR | PRAME_16 | 5 | sp\|Q92560\|BAP1_HUMAN | BAP1_5 |
| 6 | tr\|A0A2K6LQQ8\|PRAME_RHIBE | PRAME_6 | 17 | tr\|A0A2R8ZVA2\|PRAME_PANPA | PRAME_17 | 6 | sp\|D3ZHS6\|BAP1_RAT | BAP1_6 |
| 7 | tr\|A0A2K5J4S6\|PRAME_COLAP | PRAME_7 | 18 | sp\|P78395\|PRAME_HUMAN | PRAME_18 | 7 | sp\|Q5F3N6\|BAP1_CHICK | BAP1_7 |
| 8 | tr\|A0A8C9GCL8\|PRAME_9PRIM | PRAME_8 | 19 | tr\|G7PHC8\|PRAME_MACFA | PRAME_19 | 8 | sp\|A1L2G3\|BAP1_DANRE | BAP1_8 |
| 9 | tr\|A0A0D9RN10\|PRAME_CHLSB | PRAME_9 | 20 | tr\|A0A8I5YKE1\|PRAME_PONAB | PRAME_20 | 9 | sp\|C4A0D9\|BAP1_BRAFL | BAP1_9 |
| 10 | tr\|G3R981\|PRAME_GORGO | PRAME_10 | 21 | tr\|G3RVX9\|PRAME_GORGO | PRAME_21 |  |  |  |
| 11 | tr\|A0A2J8RZ57\|PRAME_PONAB | PRAME_11 |  |  |  |  |  |  |

Furthermore, we have listed organisms associated to each unique PRAME and BAP1 sequences as mentioned in Table 2. Representative images of the organisms associated to these PRAME and BAP1 sequences were presented in Figures 1 and 2

**Table 2:**
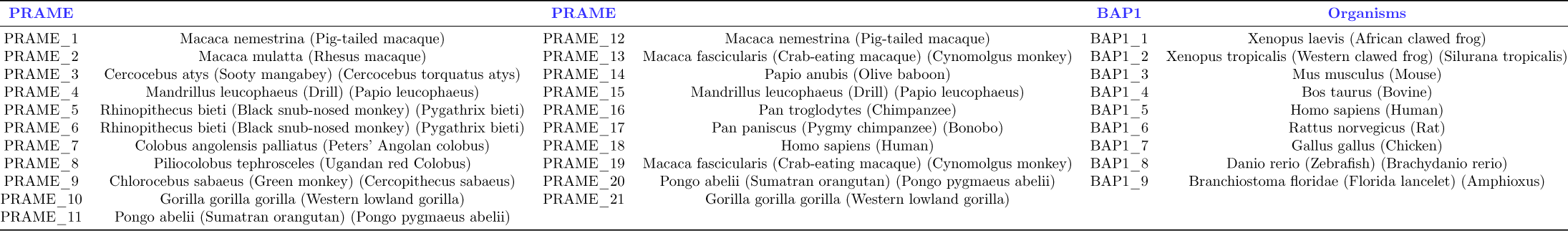
List of organisms for each PRAME and BAP1 sequence.

**Figure 1:**
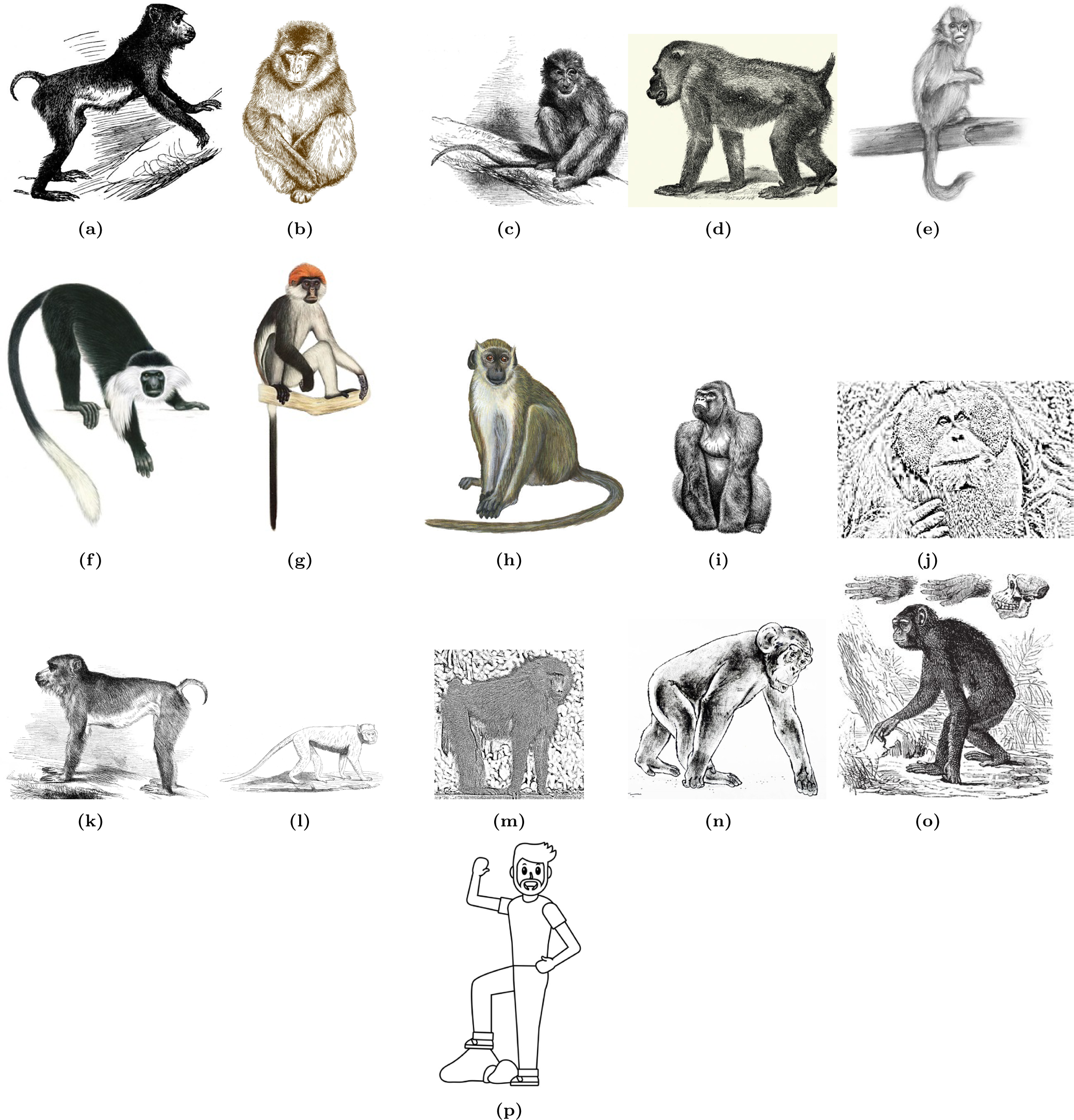
Organisms associated with PRAME proteins: a) Pig-tailed macaque, b) Rhesus macaque, c) Sooty mangabey, d) Drill, e) Black snub-nosed monkey, f) Peters’ Angolan colobus, g) Ugandan red Colobus, h) Green monkey, i) Western lowland gorilla, j) Pongo pygmaeus abelii, k) Pig-tailed macaque, l) Cynomolgus monkey, m) Olive baboon, n) Chimpanzee, o) Pygmy chimpanzee, p) Homo sapiens

**Figure 2:**
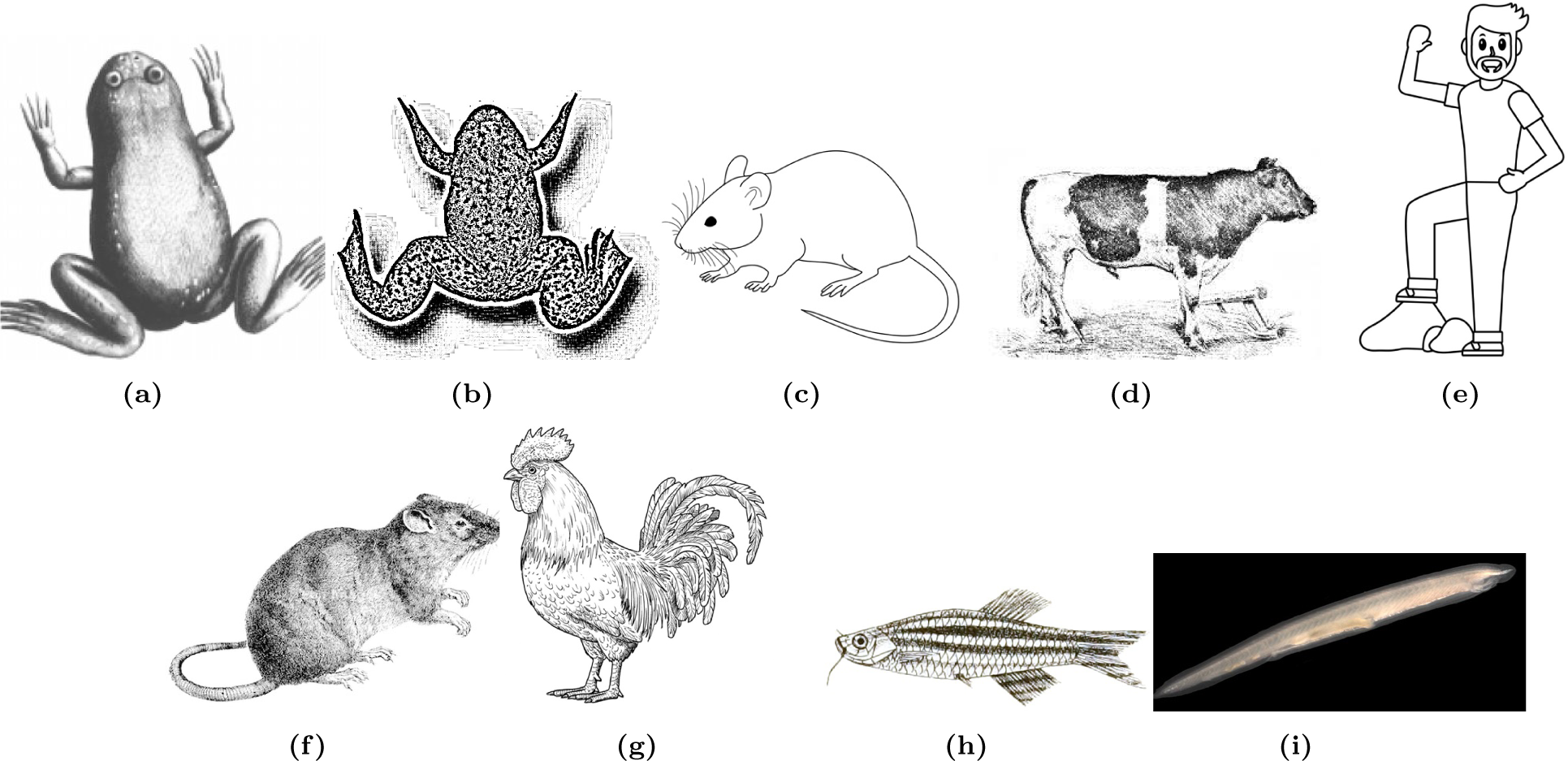
Organisms associated with BAP1 proteins: a) African clawed frog, b) Western clawed frog, c) Mus musculus, d) Bos taurus, e) Homo sapiens, f) Rattus norvegicus, g) Gallus gallus, h) Zebrafish, i) Florida lancelet.

## 3 Methods

### 3.1. Amino acid sequence homology-based phylogeny and variability of PRAME and BAP1 sequences

#### 3.1.1. Amino acid sequence homology and phylogeny

Homology analysis of 21 PRAME and 9 BAP1 sequences were made by determining multiple amino acid sequence homology, derived from Clustal Omega to identify conserved regions and variations [14, 15]. Pairwise sequence identity calculations were then employed to quantitatively assess the similarity between sequences [16]. The resulting phylogenetic tree was visualized using MATLAB phytree function, offering a graphical representation of the evolutionary relationships among the proteins [17]. This approach allowed for the interpretation of the tree in the context of sequence homology, providing insights into the parental connections and divergence events among the PRAME/BAP1 proteins [18].

#### 3.1.2. Amino acid frequency-based Shannon variability for each residue of PRAME/BAP1 sequences

Shannon entropy serves as a pivotal tool for evaluating the diversity of amino acid residues at individual positions within the alignment of PRAME (BAP1) sequences. The Shannon variability (*H_v_*) is utilized to quantify the extent of variability at each position, defined by the equation:

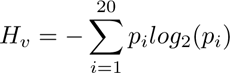

Here, *p_i_*represents the fraction of residues belonging to the amino acid type *i* at a specific position. The resulting Shannon variability (*H_v_*) ranges from 0 (indicating the presence of only one residue at that position) to 4.322 (indicating an equal distribution of all 20 amino acid residues at that position) [19]. Generally, positions with *H_v_ >* 2.0 are deemed *variable*, while those with *H_v_ <* 2.0 are considered *conserved*. Positions with *H_v_ <* 1.0 are specifically labeled as *highly conserved* [19].

The analysis of per-residue variability through Shannon entropy offers valuable insights into functionally significant residues within the receptor protein family. Furthermore, this approach aids in the identification of potential drug targets within the receptor protein family. Residues exhibiting high variability among sequences may suggest increased flexibility and adaptability in their binding pockets, rendering them more receptive to small molecule binding [20].

### 3.2. Composition profiler of PRAME/BAP1 sequences

We utilized the Composition Profiler tool to create amino acid composition profiles for all the PRAME and BAP1 proteins examined in this study, as described in the work by Vacic et al. (2007) [21]. In this analysis, the set of amino acid sequences under investigation served as the query set, while the ‘Protein Data Bank Select 25’ was used as the background set.

Additionally, we generated a composition profile for experimentally verified disordered proteins, sourced from the DisProt database. These profiles present plots that illustrate the normalized enrichment or depletion of a specific amino acid residue, calculated using the formula 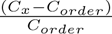, where *C_x_* represents the content of a given residue in the query protein, and *C_order_* signifies the content of the same residue in the PDB Select 25 [22, 23, 24].

### 3.3. Determining amino acid frequency composition of PRAME/BAP1 sequences

The amino acid frequency, which refers to the count of each individual amino acid within a given sequence, was calculated for all PRAME and BAP1 proteins, as discussed in previous studies [25, 26, 27]. Additionally, the relative frequency of amino acids in a sequence was determined by dividing the amino acid frequencies by the length of the respective sequence and multiplying the result by 100.

Consequently, the relative frequency of the 20 amino acids in each sequence can be visualized as a 20-dimensional vector, representing the composition of amino acids within each protein sequence.

#### 3.3.1. Evaluating Shannon entropy of PRAME/BAP1 sequences

Shannon entropy (SE) is a measure of the information content in a system [28, 29]. SE of each protein sequence is evaluated by the formula:

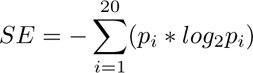

*p_i_* is the count of amino acid *i* in that sequence divided by the length of that sequence [30, 31]. *SE* reflects the degree of randomness in the amino acids count in a given sequence. A higher value of *SE* indicates greater diversity. The value of SE ranges between 0 and 4.32.

### 3.4. Determining homogeneous poly-string frequency of amino acids in PRAME/BAP1 sequences

A homogeneous poly-string of length *n* is characterized as a sequence consisting of *n* consecutive instances of a specific amino acid, as defined by Nawn et al. [32]. For example, in the sequence ‘KKKLLKKLL’, we can identify one homogeneous poly-string of K with a length of 3, one with a length of 2, and two homogeneous poly-strings of L with a length of 2. It is important to emphasize that when tallying homogeneous poly-strings of length *n*, only the exclusive and precise occurrences of that length *n* are considered.

To ascertain the maximum length of homogeneous poly-strings for all amino acids across all sequences, we tabulated the counts of homogeneous poly-strings for all conceivable lengths, ranging from 1 to the maximum length, for each amino acid within a given protein sequence.

### 3.5. Evaluating polar, non-polar residue profiles of PRAME/BAP1 sequences

In a given protein sequence (PRAME/BAP1), every amino acid was classified as either polar (represented as P) or non-polar (represented as Q). Subsequently, the protein sequence underwent transformation into a binary sequence using two symbols: P and Q. This binary P-Q profile served the purpose of representing the spatial arrangement of polar and non-polar residues within the protein sequence [32, 33, 34].

#### 3.5.1. Change response sequences based on polar, non-polar residue profiles

There are four possible transitions between two consecutive residues in a polar and non-polar profile: Polar to Polar (PP), Polar to Non-polar (PN), Non-polar to Non-polar (NN), and Non-polar to Polar (NP), as outlined in the study by Nawn et al. [32]. These transitions were documented in the form of a sequence, considering the sequential spatial arrangement of polar and non-polar residues within a given P-Q binary profile. This sequence is referred to as the “P-Q Change Response Sequence (*CRS_P Q_*)”.

The occurrence frequency of each of these four types of changes was tabulated from the binary polar and non-polar profiles corresponding to each PRAME/BAP1 protein.

### 3.6. Evaluating acidic, basic, neutral residue profiles of PRAME/BAP1 sequences

Every amino acid in a given PRAME/BAP1 protein sequence was classified as either acidic (represented as A), basic (represented as B), or neutral (represented as N). As a result, each protein sequence underwent transformation into a ternary-valued sequence, represented by A-B-N profiles, utilizing three distinct symbols: A, B, and N.

#### 3.6.1. Change response sequences based on acidic-basic-neutral residue profiles

There are nine possible transitions between two consecutive residues in an A-B-N profile: Acidic to Acidic (AA), Acidic to Basic (AB), Acidic to Neutral (AN), Basic to Acidic (BA), Basic to Basic (BB), Basic to Neutral (BN), Neutral to Acidic (NA), Neutral to Basic (NB), and Neutral to Neutral (NN). These transitions were documented in the form of a sequence, considering the sequential spatial arrangement of acidic, basic, and neutral residues within a given A-B-N ternary profile. This sequence is referred to as the “A-B-N Change Response Sequence (*CRS_ABN_*)” [32].

The frequency of each of these nine types of changes was tabulated for a given ternary A-B-N profile corresponding to each PRAME/BAP1 protein.

### 3.7. Evaluating intrinsic protein disorder

In our examination of PRAME and BAP1 proteins in this study, we evaluated their propensity for intrinsic disorder using a collection of widely utilized per-residue disorder prediction tools, including PONDR® VLS2, PONDR® VL3, PONDR® VLXT, PONDR® FIT, IUPred-Long, and IUPred-Short, as referenced in the literature [35, 36, 37, 38, 39, 40]. To facilitate this analysis, we utilized the Rapid Intrinsic Disorder Analysis Online (RIDAO) web platform to aggregate results from each predictor comprehensively [41].The percentage of predicted intrinsically disordered residues (PPIDR) for each protein served as the basis for their classification according to the degree of disorder. [42, 43].

The per-residue disorder score ranged from 0 to 1, where a score of 0 denoted fully ordered residues and a score of 1 indicated fully disordered residues. Residues with scores surpassing the threshold of 0.5 were categorized as “disordered residues” (‘D’). Those with disorder scores between 0.25 and 0.5 were deemed “highly flexible” (‘HF’), while those with scores between 0.1 and 0.25 were classified as “moderately flexible” (‘MF’), in accordance with the methodology outlined in the literature [40]. Other residues (disorder scores less than 0.1) were denoted as ‘O’.

#### 3.7.1. Percentage of four intrinsic protein disorder residue types in each amino acid

As the distributions of amino acids were non-uniform (relative frequency of some amino acids were very small amount while some had very large frequency), counts of each of the four residue types (‘D’, ‘HF’, ‘MF’ and ‘O’) for a particular amino acid were divided by the corresponding amino acid frequency and multiplied by 100 in each sequence. It is to be noted that percentages were calculated based on individual amino acid frequency and not with respect to the total number of residues of a given type in a sequence. Hence the sum of the percentages of different amino acids will not be 100 for a given residue type in a sequence. Even if the frequency of an amino acid is very less in any sequence, its distribution among four residue types can be better reflected in this way.

#### 3.7.2. Change response sequences based on intrinsic protein disorder residues

There are sixteen potential transitions between consecutive residues in PRAME and BAP1 protein sequences. These transitions encompass disordered to disordered (D_D), disordered to highly flexible (D_HF), disordered to moderately flexible (D_MF), disordered to other (D_O), and the corresponding twelve transitions (HF_D, HF_HF, HF_MF, HF_O, MF_D, MF_HF, MF_MF, MF_O, O_D, O_HF, O_MF, and O_O). These transitions were documented in the form of a sequence, considering the sequential spatial arrangement of D, HF, MF, and O residues within each sequence. This sequence is referred to as the “Change Response Sequence,” and the frequency of each of the sixteen changes was tabulated from these sequences, as outlined in the work by Nawn et al. [32].

### 3.8. Phylogenetic relationships among PRAME/BAP1 proteins

We assessed the Euclidean distance between feature vectors for all pairs of PRAME/BAP1 protein sequences based on four features namely, relative frequency of amino acids, relative frequency of four changes obtained from polar-nonpolar residues, the relative frequency of nine changes obtained from acidic-basic-neutral residues, the relative frequency of sixteen changes obtained from intrinsic protein disorder analysis [32, 44]. Each feature generated a distance matrix of dimensions 21 × 21 (for PRAME)/9 × 9 (for BAP1). Dendrograms (phylogenies) were constructed based on Euclidean distances and average linkage.

Sequence homology-based similarity matrices were obtained from Clustal Omega. Distance matrices for sequence homology were obtained by subtracting similarity matrices from 100.

#### 3.8.1. Proximal set of PRAME/BAP1 proteins

Proximity of PRAME/BAP1 sequences were derived from the phylogenetic relationships at different scales. It is to be noted that the scale level was not equal to the distance value in y axis of dendrograms. Adjacent sequences (sequences which were merged) below a certain distance level in the dendrograms were termed ‘proximal sequences’ and the set they formed was called ‘proximal set’ at that scale level. Different ‘proximal sets’ at a lower scale level may be merged to form one‘proximal set’ at upper scale.

## 4. Results and analyses

The initial step in characterizing PRAME/BAP1 proteins from diverse organisms involves utilizing multiple sequence alignment (MSA) to assess sequence homology in a quantitative manner. We first analyzed distances between all pairs of PRAME/BAP1 sequences, based on dissimilarity derived from MSA. Subsequently, the quantification of amino acid variability per-residue position for each sequence was conducted using Shannon entropy. Then we investigated amino acid compositions and the relative percentage of amino acids present in PRAME/BAP1 sequences. In addition, we also studied the frequency of homogeneous poly-strings of amino acids present in PRAME/BAP1 sequences. Furthermore, polar and non-polar residue compositions and acidic, basic, and neutral residue compositions of amino acid sequences of PRAME/BAP1 were studied to develop phylogenetic relationships. Intrinsic protein disorder analysis of PRAME and BAP1 proteins was also carried out.

### 4.1. Sequence homology, invariant residues, and per-residue Shannon variability

In this section, phylogenetic relationships for PRAME and BAP1 sequences were developed based on amino acid sequence homology derived from Clustal Omega. Subsequently, per-residue amino acid variability was enumerated using Shannon entropy. Furthermore, invariant residues and their impact were discussed.

#### 4.1.1. Sequence homology based phylogenetic relationships

Figure 3 illustrates the phylogenetic relationship between the PRAME (top) and BAP1 (bottom) proteins, determined by sequence homology. Notably, PRAME_10 and PRAME_21 exhibited 100% sequence homology as indicated by the PRAME sequence phylogeny, despite their original non-identical status. PRAME_10 (with a length of 509) and PRAME_21 (with a length of 423), originating from the same organism, the Western lowland gorilla, share a common ancestral signature-sequence, highlighted in violet in Figure 4. Three proximal sets of PRAME sequences, denoted as {1,2}, {11,20}, and {12,13} were obtained at scale 1(Table 3). Consequently, other proximal sets were presented in different scales (in increasing scale number). It is important to highlight that each of the 21 PRAME sequences was assigned to one of these proximal sets within scale 7. In contrast, BAP1_8 and BAP1_9 were not included in any proximal set till the considered scale (refer to Table 3). Notably, the sequence BAP1_9 could be considered as an outlier, as illustrated in Figure 3.

**Figure 3:**
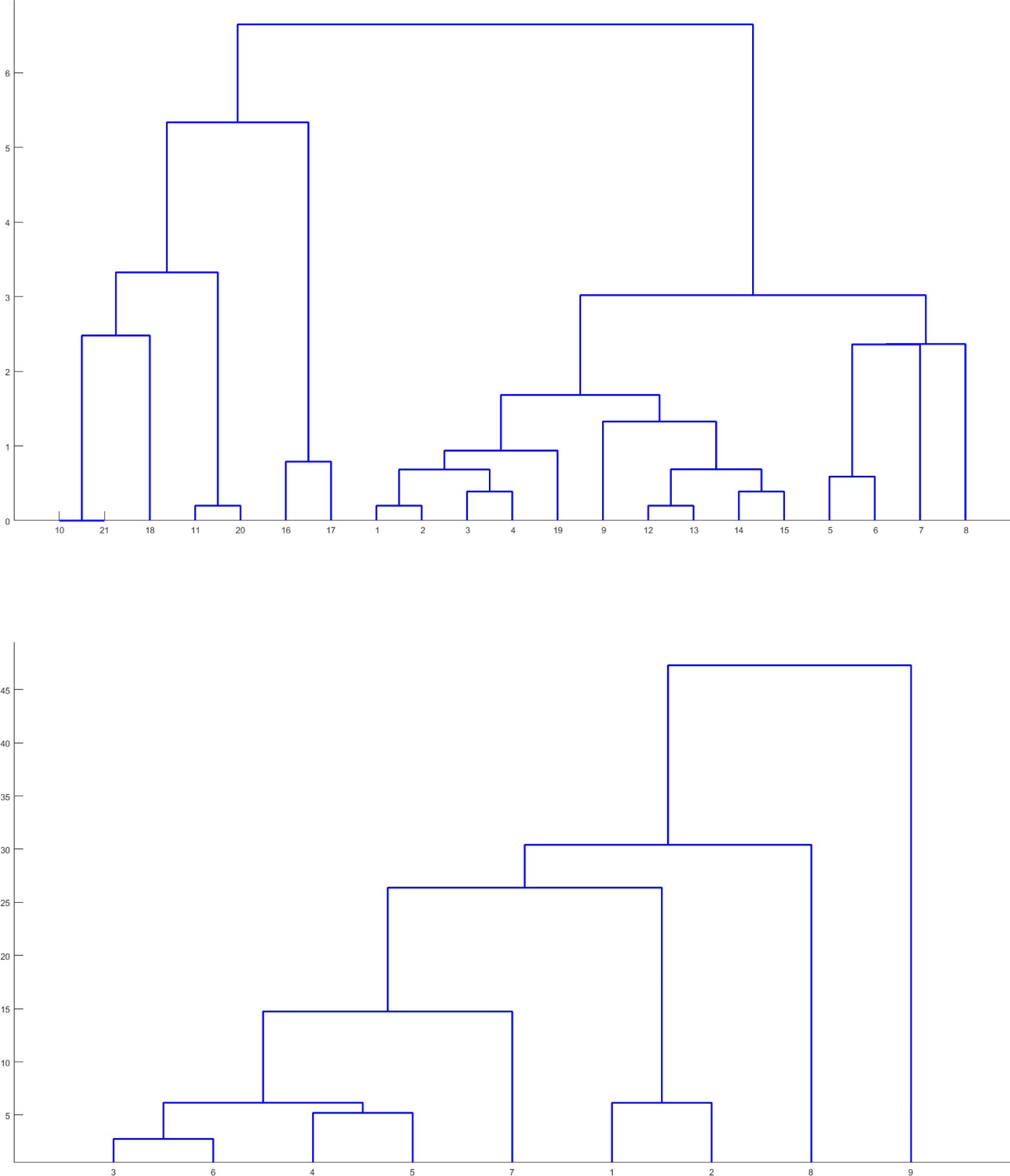
Phylogenetic relationship among the PRAME (top) and BAP1 (bottom) proteins based on sequence homology.

**Figure 4:**
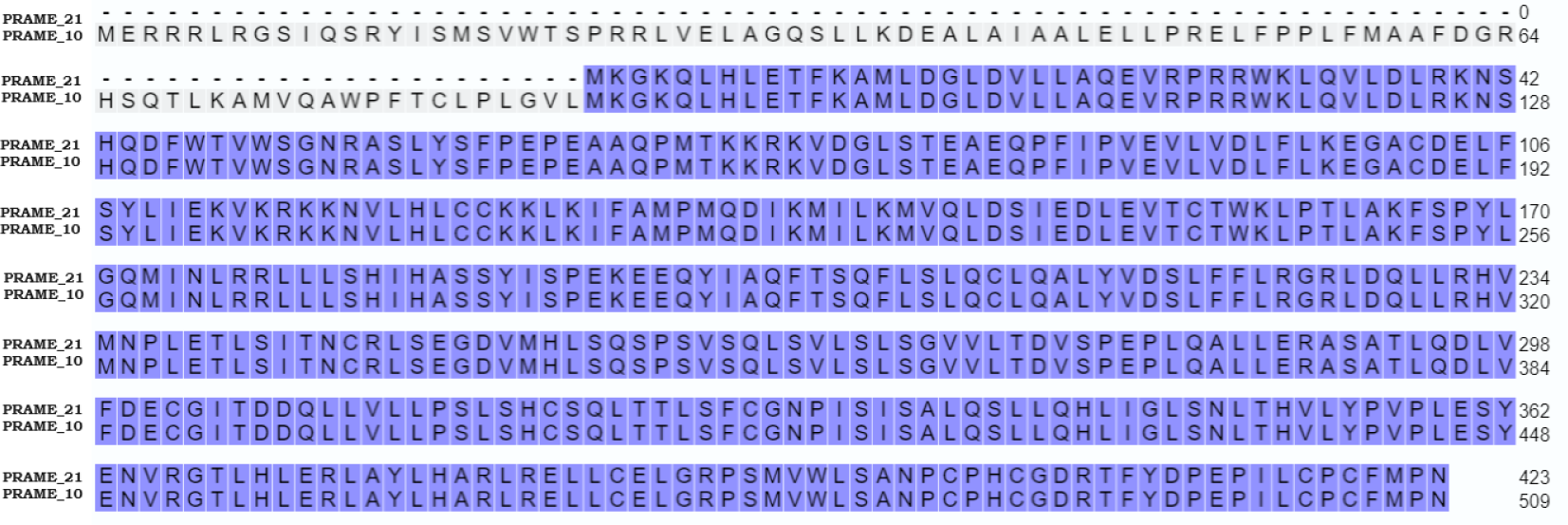
(Top): Amino acid sequence homology between PRAME_10 and PRAME_21.

**Table 3:** Scaled-based proximal sets of PRAME/BAP1 proteins based on MSA.

| *Scales* | *PRAME* | *Scales* | *BAP1* |
| --- | --- | --- | --- |
| Scale-0 | {10, 21} | Scale-0 | {3, 6} |
| Scale-1 | {1, 2}; {11, 20}; {12, 13} | Scale-1 | {{3, 6}, {4, 5}}; {1, 2} |
| Scale-2 | {14, 15}; {3, 4} | Scale-2 | {{{3, 6}, {4, 5}}, {7}} |
| Scale-3 | {5, 6} |  |  |
| Scale-4 | {16, 17}; {{1, 2}, {3, 4}}; {{12, 13}, {14, 15}} |  |  |
| Scale-5 | {{{1, 2}, {3, 4}}, {19}} |  |  |
| Scale-6 | {{{12, 13}, {14, 15}}, {9}} |  |  |
| Scale-7 | {{{5, 6}, {7}}, {8}}; {{10, 21}, {18}}; {{{1, 2}, {3, 4}}, {19}}; {{{12, 13}, {14, 15}}, {9}} |  |  |

#### 4.1.2. Shannon variability of each amino acid residue position

It was found that most of the residues in PRAME variants are highly conserved, with an impressive percentage of 97.45% (Figure 4). Similarly, in BAP1 variants, a substantial 71.55% of residues exhibit high conservation. The comparatively lower percentages of conserved residues in PRAME (2.54%) and BAP1 (22.12%) suggest that a smaller proportion of residues in these proteins show low variations in their amino acid composition [32]. The absence of variable residues in PRAME and less than 10% in BAP1 indicate a high degree of conservation in both proteins. This conservation might be indicative of important structural or functional roles for these conserved residues in maintaining the integrity or activity of these proteins across different variants. Overall, the results suggest a strong selective pressure favoring the preservation of specific amino acid residues in these proteins.

Additionally, Table 4 displays a compilation of invariant residues, characterized by a Shannon variability index of 0. Across the 21 PRAME sequences, the total number of invariant residues or amino acid positions amounted to 358, while for the 9 BAP1 sequences, it was 236 (Figure 5). Notably, 71.8% of amino residues within the highly conserved regions exhibited invariance in PRAME sequences, whereas in BAP1 sequences, this percentage was 41.7%. This suggests that a significant proportion of the highly conserved regions in both PRAME and BAP1 sequences exhibit sequence invariance, highlighting the potential functional importance or evolutionary conservation of these specific amino acid positions in the studied proteins.

**Table 4:**
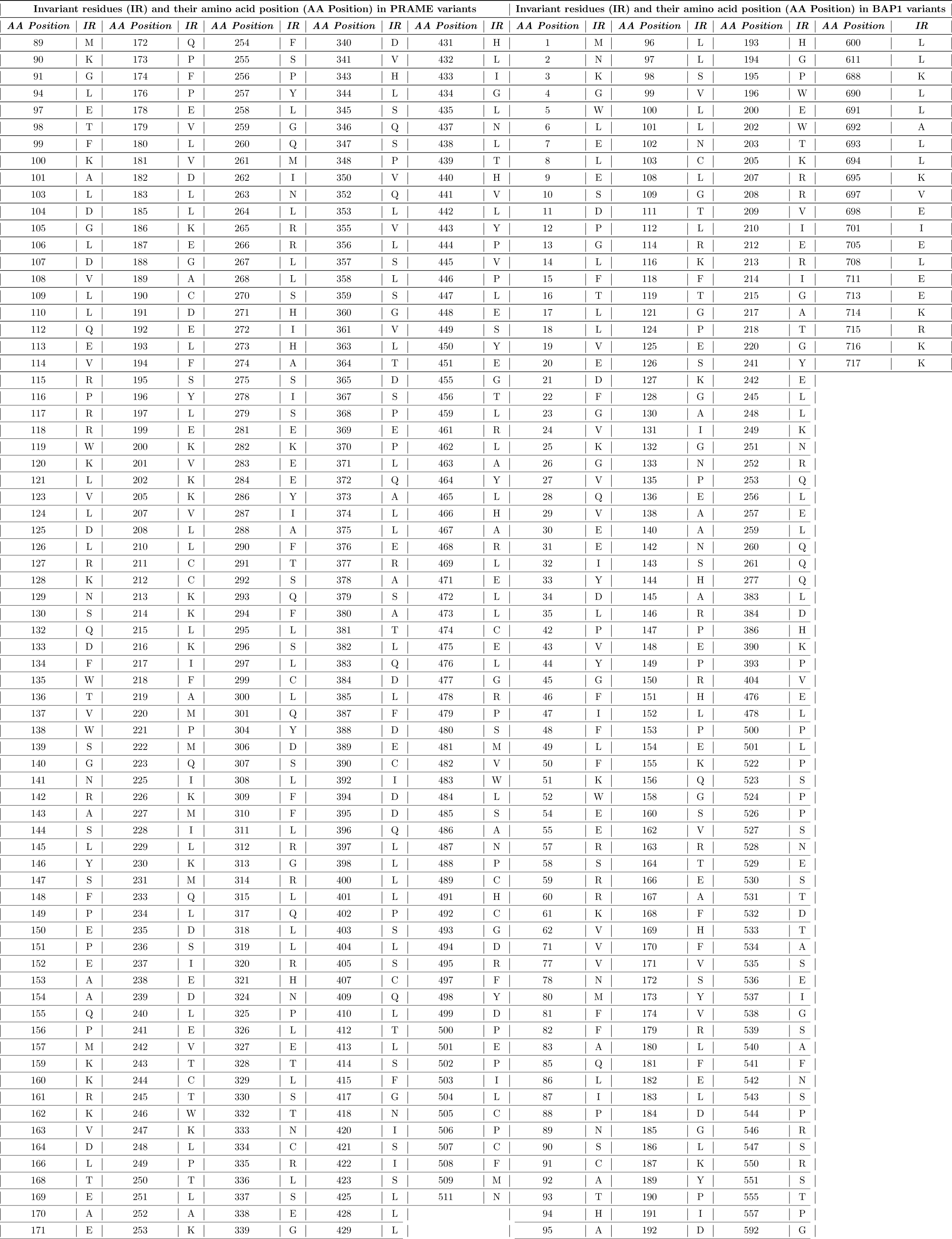
Invariant residues (IR) and their amino acid position (AA Position) in PRAME/BAP1 variants.

| Invariant residues (IR) and their amino acid position (AA Position) in PRAME variants |  |  |  |  |  |  |  |  |  | Invariant residues (IR) and their amino acid position (AA Position) in BAP1 variants |  |  |  |  |  |  |  |
| --- | --- | --- | --- | --- | --- | --- | --- | --- | --- | --- | --- | --- | --- | --- | --- | --- | --- |
| AA Position | IR | AA Position | IR | AA Position | IR | AA Position | IR | AA Position | IR | AA Position | IR | AA Position | IR | AA Position | IR | AA Position | IR |
| 89 | M | 172 | Q | 254 | F | 340 | D | 431 | H | 1 | M | 96 | L | 193 | H | 600 | L |
| 90 | K | 173 | P | 255 | S | 341 | V | 432 | L | 2 | N | 97 | L | 194 | G | 611 | L |
| 91 | G | 174 | F | 256 | P | 343 | H | 433 | I | 3 | K | 98 | S | 195 | P | 688 | K |
| 94 | L | 176 | P | 257 | Y | 344 | L | 434 | G | 4 | G | 99 | V | 196 | W | 690 | L |
| 97 | E | 178 | E | 258 | L | 345 | S | 435 | L | 5 | W | 100 | L | 200 | E | 691 | L |
| 98 | T | 179 | V | 259 | G | 346 | Q | 437 | N | 6 | L | 101 | L | 202 | W | 692 | A |
| 99 | F | 180 | L | 260 | Q | 347 | S | 438 | L | 7 | E | 102 | N | 203 | T | 693 | L |
| 100 | K | 181 | V | 261 | M | 348 | P | 439 | T | 8 | L | 103 | C | 205 | K | 694 | L |
| 101 | A | 182 | D | 262 | I | 350 | V | 440 | H | 9 | E | 108 | L | 207 | R | 695 | K |
| 103 | L | 183 | L | 263 | N | 352 | Q | 441 | V | 10 | S | 109 | G | 208 | R | 697 | V |
| 104 | D | 185 | L | 264 | L | 353 | L | 442 | L | 11 | D | 111 | T | 209 | V | 698 | E |
| 105 | G | 186 | K | 265 | R | 355 | V | 443 | Y | 12 | P | 112 | L | 210 | I | 701 | I |
| 106 | L | 187 | E | 266 | R | 356 | L | 444 | P | 13 | G | 114 | R | 212 | E | 705 | E |
| 107 | D | 188 | G | 267 | L | 357 | S | 445 | V | 14 | L | 116 | K | 213 | R | 708 | L |
| 108 | V | 189 | A | 268 | L | 358 | L | 446 | P | 15 | F | 118 | F | 214 | I | 711 | E |
| 109 | L | 190 | C | 270 | S | 359 | S | 447 | L | 16 | T | 119 | T | 215 | G | 713 | E |
| 110 | L | 191 | D | 271 | H | 360 | G | 448 | E | 17 | L | 121 | G | 217 | A | 714 | K |
| 112 | Q | 192 | E | 272 | I | 361 | V | 449 | S | 18 | L | 124 | P | 218 | T | 715 | R |
| 113 | E | 193 | L | 273 | H | 363 | L | 450 | Y | 19 | V | 125 | E | 220 | G | 716 | K |
| 114 | V | 194 | F | 274 | A | 364 | T | 451 | E | 20 | E | 126 | S | 241 | Y | 717 | K |
| 115 | R | 195 | S | 275 | S | 365 | D | 455 | G | 21 | D | 127 | K | 242 | E |  |  |
| 116 | P | 196 | Y | 278 | I | 367 | S | 456 | T | 22 | F | 128 | G | 245 | L |  |  |
| 117 | R | 197 | L | 279 | S | 368 | P | 459 | L | 23 | G | 130 | A | 248 | L |  |  |
| 118 | R | 199 | E | 281 | E | 369 | E | 461 | R | 24 | V | 131 | I | 249 | K |  |  |
| 119 | W | 200 | K | 282 | K | 370 | P | 462 | L | 25 | K | 132 | G | 251 | N |  |  |
| 120 | K | 201 | V | 283 | E | 371 | L | 463 | A | 26 | G | 133 | N | 252 | R |  |  |
| 121 | L | 202 | K | 284 | E | 372 | Q | 464 | Y | 27 | V | 135 | P | 253 | Q |  |  |
| 123 | V | 205 | K | 286 | Y | 373 | A | 465 | L | 28 | Q | 136 | E | 256 | L |  |  |
| 124 | L | 207 | V | 287 | I | 374 | L | 466 | H | 29 | V | 138 | A | 257 | E |  |  |
| 125 | D | 208 | L | 288 | A | 375 | L | 467 | A | 30 | E | 140 | A | 259 | L |  |  |
| 126 | L | 210 | L | 290 | F | 376 | E | 468 | R | 31 | E | 142 | N | 260 | Q |  |  |
| 127 | R | 211 | C | 291 | T | 377 | R | 469 | L | 32 | I | 143 | S | 261 | Q |  |  |
| 128 | K | 212 | C | 292 | S | 378 | A | 471 | E | 33 | Y | 144 | H | 277 | Q |  |  |
| 129 | N | 213 | K | 293 | Q | 379 | S | 472 | L | 34 | D | 145 | A | 383 | L |  |  |
| 130 | S | 214 | K | 294 | F | 380 | A | 473 | L | 35 | L | 146 | R | 384 | D |  |  |
| 132 | Q | 215 | L | 295 | L | 381 | T | 474 | C | 42 | P | 147 | P | 386 | H |  |  |
| 133 | D | 216 | K | 296 | S | 382 | L | 475 | E | 43 | V | 148 | E | 390 | K |  |  |
| 134 | F | 217 | I | 297 | L | 383 | Q | 476 | L | 44 | Y | 149 | P | 393 | P |  |  |
| 135 | W | 218 | F | 299 | C | 384 | D | 477 | G | 45 | G | 150 | R | 404 | V |  |  |
| 136 | T | 219 | A | 300 | L | 385 | L | 478 | R | 46 | F | 151 | H | 476 | E |  |  |
| 137 | V | 220 | M | 301 | Q | 387 | F | 479 | P | 47 | I | 152 | L | 478 | L |  |  |
| 138 | W | 221 | P | 304 | Y | 388 | D | 480 | S | 48 | F | 153 | P | 500 | P |  |  |
| 139 | S | 222 | M | 306 | D | 389 | E | 481 | M | 49 | L | 154 | E | 501 | L |  |  |
| 140 | G | 223 | Q | 307 | S | 390 | C | 482 | V | 50 | F | 155 | K | 522 | P |  |  |
| 141 | N | 225 | I | 308 | L | 392 | I | 483 | W | 51 | K | 156 | Q | 523 | S |  |  |
| 142 | R | 226 | K | 309 | F | 394 | D | 484 | L | 52 | W | 158 | G | 524 | P |  |  |
| 143 | A | 227 | M | 310 | F | 395 | D | 485 | S | 54 | E | 160 | S | 526 | P |  |  |
| 144 | S | 228 | I | 311 | L | 396 | Q | 486 | A | 55 | E | 162 | V | 527 | S |  |  |
| 145 | L | 229 | L | 312 | R | 397 | L | 487 | N | 57 | R | 163 | R | 528 | N |  |  |
| 146 | Y | 230 | K | 313 | G | 398 | L | 488 | P | 58 | S | 164 | T | 529 | E |  |  |
| 147 | S | 231 | M | 314 | R | 400 | L | 489 | C | 59 | R | 166 | E | 530 | S |  |  |
| 148 | F | 233 | Q | 315 | L | 401 | L | 491 | H | 60 | R | 167 | A | 531 | T |  |  |
| 149 | P | 234 | L | 317 | Q | 402 | P | 492 | C | 61 | K | 168 | F | 532 | D |  |  |
| 150 | E | 235 | D | 318 | L | 403 | S | 493 | G | 62 | V | 169 | H | 533 | T |  |  |
| 151 | P | 236 | S | 319 | L | 404 | L | 494 | D | 71 | V | 170 | F | 534 | A |  |  |
| 152 | E | 237 | I | 320 | R | 405 | S | 495 | R | 77 | V | 171 | V | 535 | S |  |  |
| 153 | A | 238 | E | 321 | H | 407 | C | 497 | F | 78 | N | 172 | S | 536 | E |  |  |
| 154 | A | 239 | D | 324 | N | 409 | Q | 498 | Y | 80 | M | 173 | Y | 537 | I |  |  |
| 155 | Q | 240 | L | 325 | P | 410 | L | 499 | D | 81 | F | 174 | V | 538 | G |  |  |
| 156 | P | 241 | E | 326 | L | 412 | T | 500 | P | 82 | F | 179 | R | 539 | S |  |  |
| 157 | M | 242 | V | 327 | E | 413 | L | 501 | E | 83 | A | 180 | L | 540 | A |  |  |
| 159 | K | 243 | T | 328 | T | 414 | S | 502 | P | 85 | Q | 181 | F | 541 | F |  |  |
| 160 | K | 244 | C | 329 | L | 415 | F | 503 | I | 86 | L | 182 | E | 542 | N |  |  |
| 161 | R | 245 | T | 330 | S | 417 | G | 504 | L | 87 | I | 183 | L | 543 | S |  |  |
| 162 | K | 246 | W | 332 | T | 418 | N | 505 | C | 88 | P | 184 | D | 544 | P |  |  |
| 163 | V | 247 | K | 333 | N | 420 | I | 506 | P | 89 | N | 185 | G | 546 | R |  |  |
| 164 | D | 248 | L | 334 | C | 421 | S | 507 | C | 90 | S | 186 | L | 547 | S |  |  |
| 166 | L | 249 | P | 335 | R | 422 | I | 508 | F | 91 | C | 187 | K | 550 | R |  |  |
| 168 | T | 250 | T | 336 | L | 423 | S | 509 | M | 92 | A | 189 | Y | 551 | S |  |  |
| 169 | E | 251 | L | 337 | S | 425 | L | 511 | N | 93 | T | 190 | P | 555 | T |  |  |
| 170 | A | 252 | A | 338 | E | 428 | L |  |  | 94 | H | 191 | I | 557 | P |  |  |
| 171 | E | 253 | K | 339 | G | 429 | L |  |  | 95 | A | 192 | D | 592 | G |  |  |

**Figure 5:**
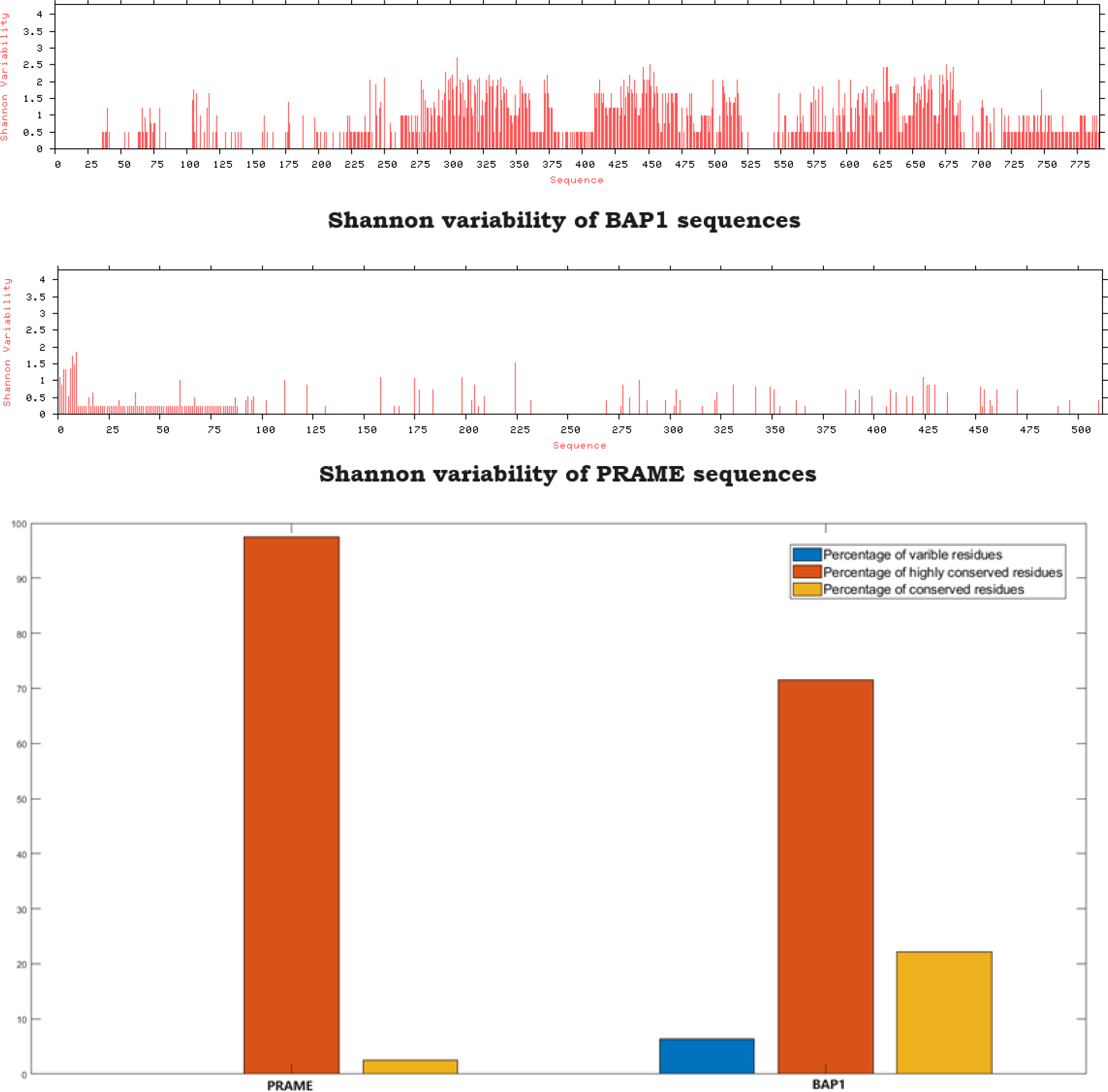
Shannon variability of amino acid residues in PRAME and BAP1 proteins.

### 4.2. Sequence-based amino acid frequency composition and Shannon entropy

#### 4.2.1. Compositional profile of PRAME and BAP1 proteins

Marked by noticeable distinctions, research has demonstrated that disordered proteins or regions exhibit a significant scarcity of bulky hydrophobic amino acid residues (I, L, and V) and aromatic amino acids (W, Y, F, and H). These latter residues often play a role in constructing the hydrophobic core of a folded globular protein. Additionally, disordered proteins or regions tend to have a low content of C, N, and M residues. The amino acids C, W, I, Y, F, L, H, V, N, and M, which are notably deficient in disordered proteins and regions, are categorized as order-promoting amino acids. Conversely, disordered proteins and regions are notably enriched in disorder-promoting amino acids, including R, T, D, G, A, K, Q, S, E, and P [21, 39, 45]. The biases in amino acid composition can be effectively visualized through a web-based tool known as the Composition Profiler. This tool enables the semi-automatic identification of amino acid enrichment or depletion in queried proteins [21].

Examining the amino acid composition profiles of 21 PRAME sequences showed a significant depletion of four order-promoting residues (I, Y, V, and M), while four disorder-promoting residues (R, Q, S, and P) were notably enriched (Figure 6).

**Figure 6:**
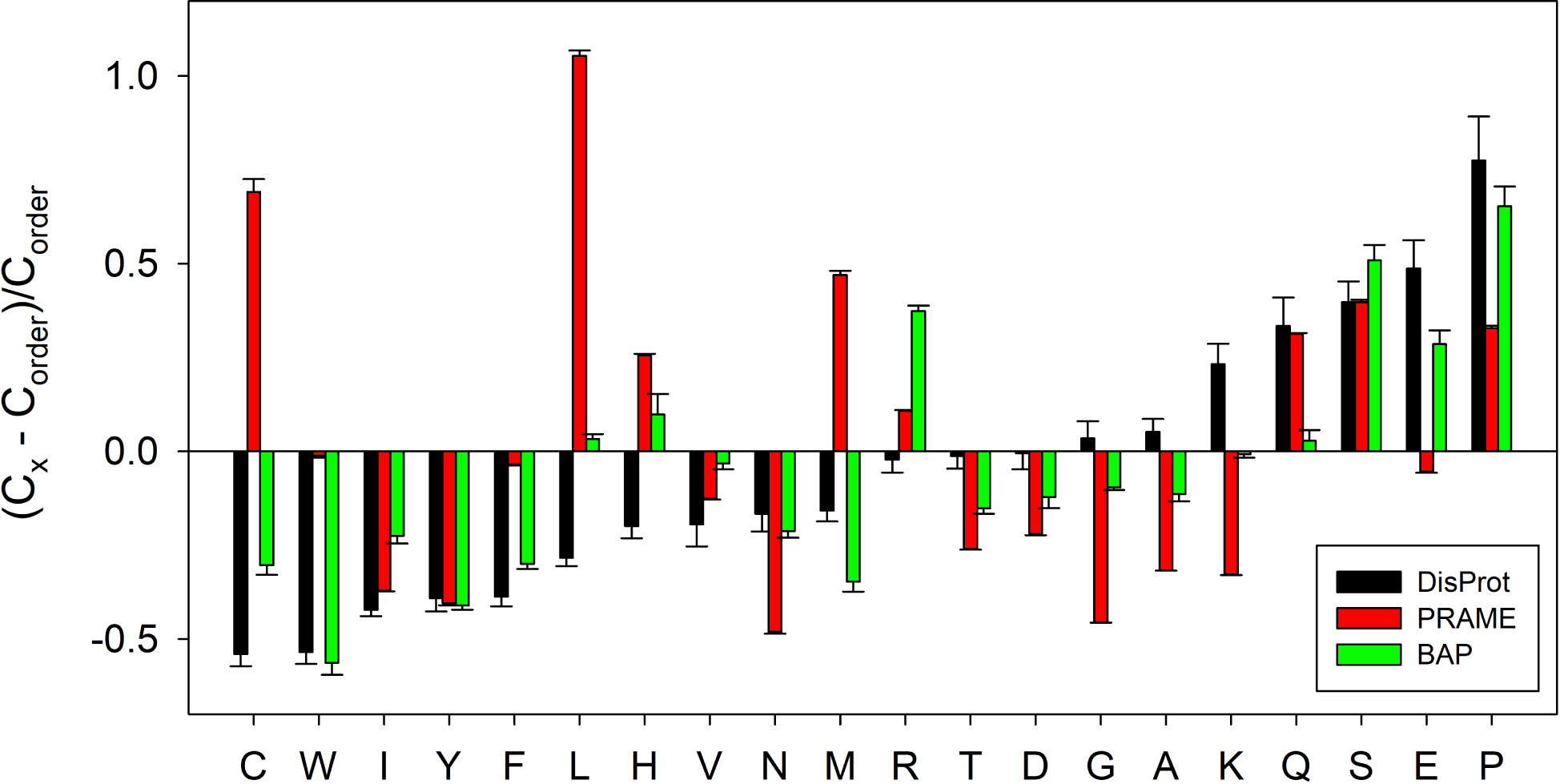
Amino acid composition profile of 21 PRAME (red bars)and 9 BAP1 proteins (green bars). The fractional difference is calculated as 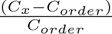, where *C_x_* is the content of a given amino acid in the query set, and *C_order_* is the content of a given amino acid in the background set (Protein Data Bank Select 25). The amino acid residues are ranked from most order-promoting residue to most disorder-promoting residue. Positive values indicate enrichment, and negative values indicate depletion of a particular amino acid. The composition profile generated for experimentally validated disordered proteins from the DisProt database (black bars) is shown for comparison. In both cases, error bars correspond to standard deviations over 10,000 bootstrap iterations.

In the analysis of amino acid composition profiles for 9 BAP1 sequences, seven order-promoting residues (C, W, I, Y, F, N, and M) were significantly depleted, while four disorder-promoting residues (R, S, E, and P) were significantly enriched (Figure 6).

The observed reduction in order-promoting residues (I, Y, V, and M) and simultaneous increase in disorder-promoting residues (R, Q, S, and P) within the amino acid composition profiles of PRAME sequences suggest a departure from the typical compositional characteristics associated with well-structured proteins. This shift in residue distribution hints at a potential inclination towards intrinsic disorder or conformational flexibility within the PRAME protein family.

Likewise, the notable decrease in order-promoting residues (C, W, I, Y, F, N, and M) and the corresponding rise in disorder-promoting residues (R, S, E, and P) in the amino acid composition profiles of BAP1 sequences indicate a distinct compositional signature. This specific pattern implies a potential predisposition towards intrinsic disorder or conformational flexibility in BAP1 proteins.

The compositional disparities between PRAME and BAP1 sequences underscore the likelihood of unique structural and functional attributes, emphasizing the role of amino acid composition as a determinant of protein behavior and characteristics. To comprehend the implications of these compositional variances in the context of PRAME and BAP1 protein functionalities, further structural and functional analyses are warranted.

#### 4.2.2. Relative frequency of amino acids and associated phylogenetic relationship

The analysis of amino acid frequencies as illustrated in Figure 7, indicates a consistent prominence of leucine, exceeding 17% in all instances of PRAME, suggesting a notable functional role. Serine was present with the second highest percentage (with values between 8 to 9%). On the other hand, percentages of three amino acids (glutamic acid, leucine, and serine) in BAP1 were between 8 to 11 in almost all BAP1 sequences. The count of tryptophan was 4 in all BAP1 except BAP1_9 which had count 8. In all PRAME and BAP1 sequences, the count of tryptophan was lowest. However, there is an appreciable disparity in the percentages of tryptophan between PRAME (1.57%) and BAP1 (0.55%) sequences of human, implying potential structural or functional distinctions. These findings underscore the distinctive amino acid compositions of PRAME and BAP1, warranting further investigation into the specific implications of leucine and tryptophan in these protein sequences.

**Figure 7:**
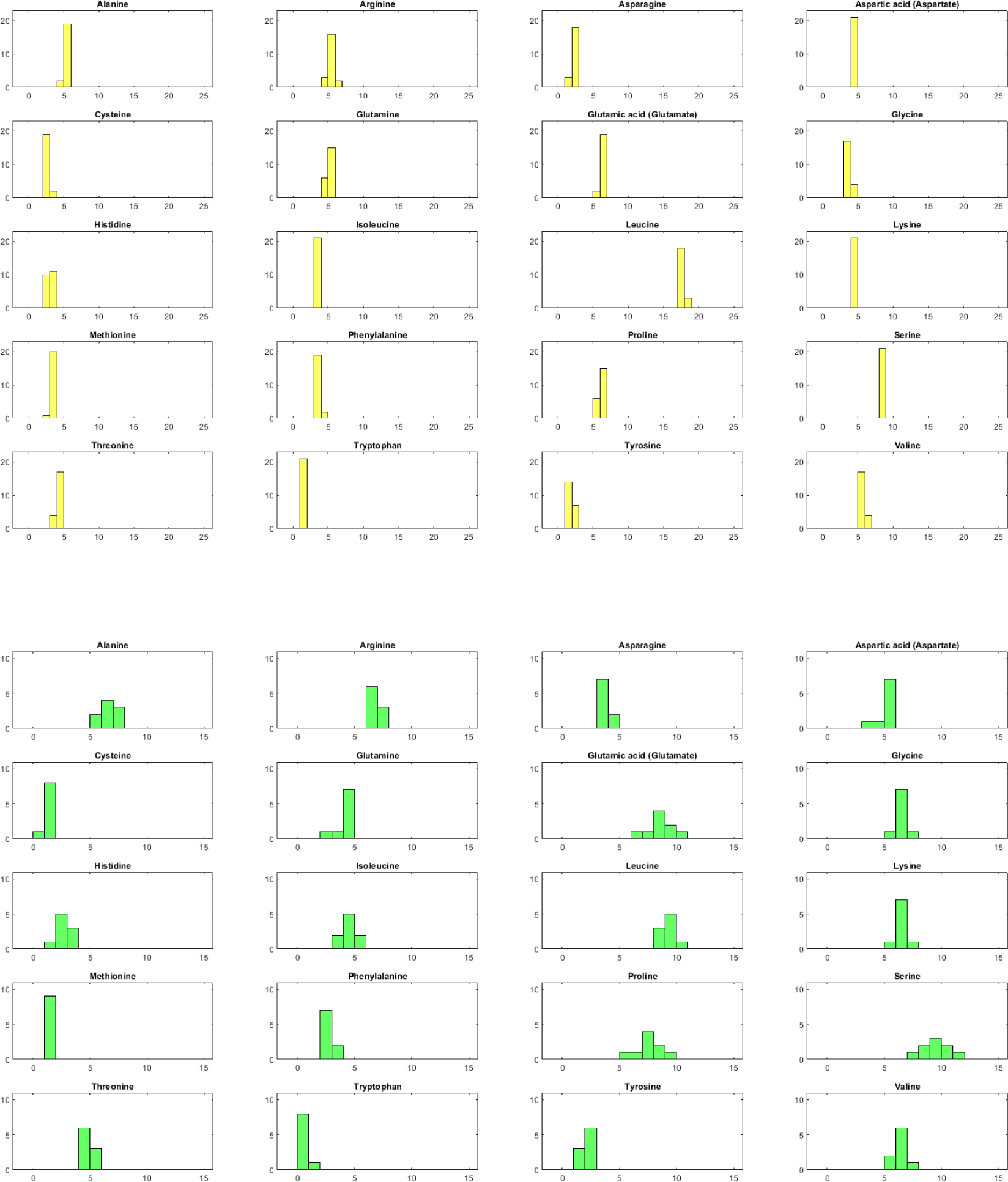
Histogram of relative frequency of each amino acid in PRAME proteins (Top), BAP1 proteins (Bottom). X axis denotes percentage and Y axis denotes number of sequences.

Figure 8 illustrates the phylogenetic relationship between PRAME (top) and BAP1 (bottom) proteins, determined by the relative frequency of each amino acid. The phylogenetic analysis identified two proximal sets labeled as {1,2} and {12,13} with corresponding scale values set at 0 (see Table 5). PRAME_21 was not included in any proximal set. For BAP1 sequences, BAP1_1 and BAP1_2 were proximal but their set was distant from the rest of the sequences (Table 5).

**Figure 8:**
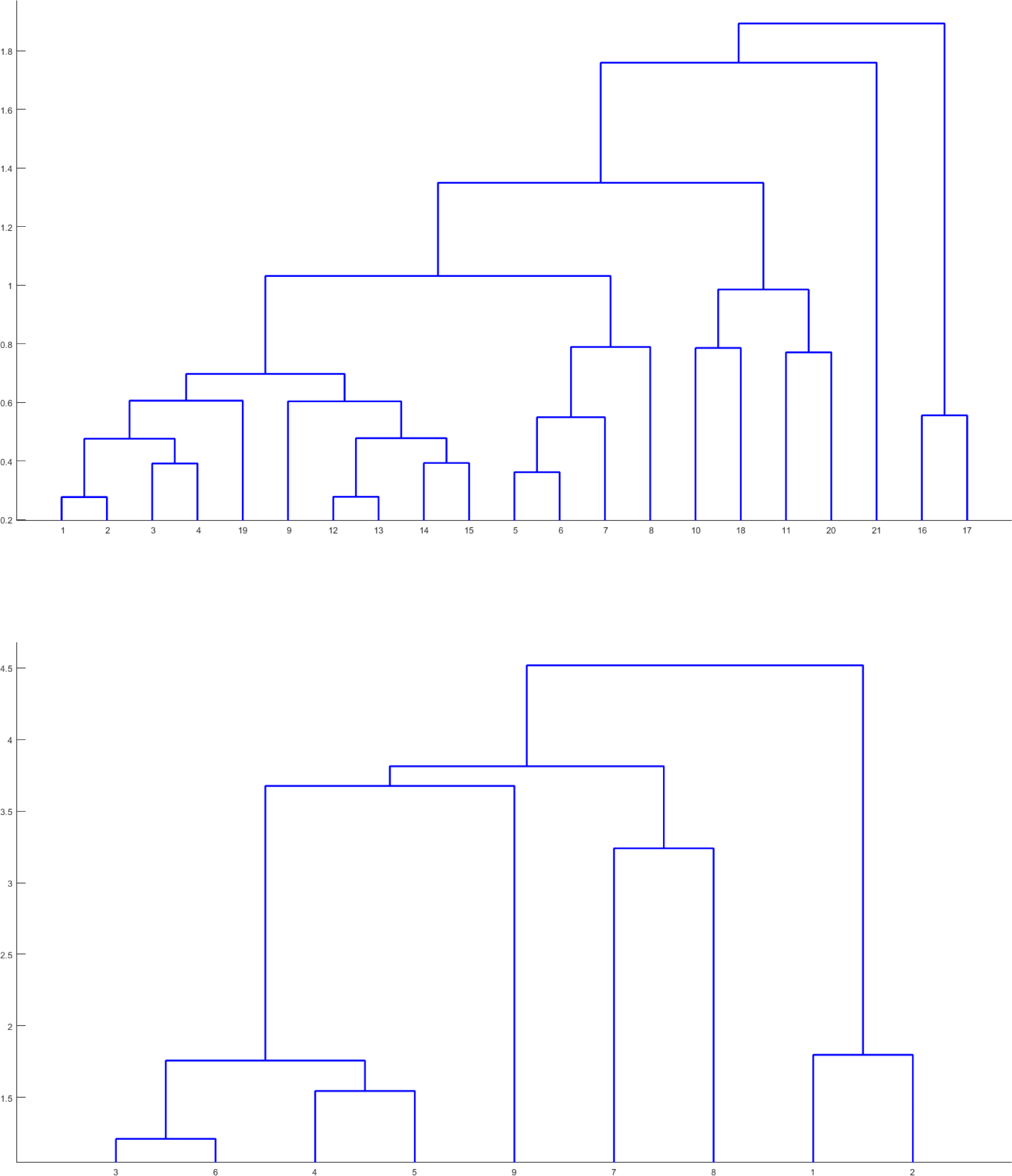
Phylogenetic relationship among the PRAME (top) and BAP1 (bottom) proteins based on relative frequency of amino acids.

**Table 5:**
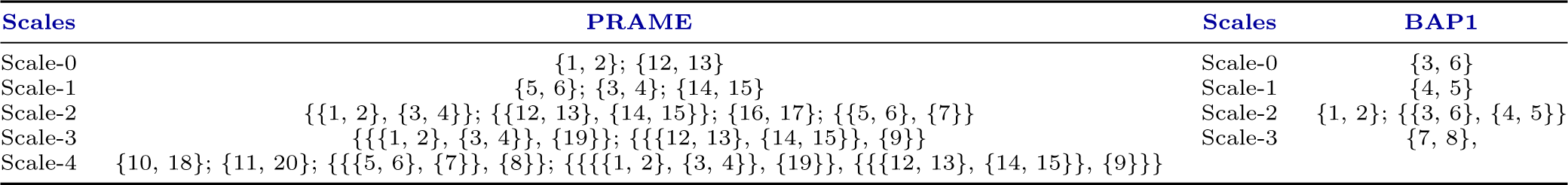
Scaled based proximal sets of PRAME/BAP1 proteins based on relative frequency of amino acids.

| Scales | PRAME | Scales | BAP1 |
| --- | --- | --- | --- |
| Scale-0 | {1, 2}; {12, 13} | Scale-0 | {3, 6} |
| Scale-1 | {5, 6}; {3, 4}; {14, 15} | Scale-1 | {4, 5} |
| Scale-2 | {{1, 2}, {3, 4}}; {{12, 13}, {14, 15}}; {16, 17}; {{5, 6}, {7}} | Scale-2 | {1, 2}; {{3, 6}, {4, 5}} |
| Scale-3 | {{{1, 2}, {3, 4}}, {19}}; {{{12, 13}, {14, 15}}, {9}} | Scale-3 | {7, 8}, |
| Scale-4 | {10, 18}; {11, 20}; {{{5, 6}, {7}}, {8}}; {{{{1, 2}, {3, 4}}, {19}}, {{{12, 13}, {14, 15}}, {9}}} |  |  |

#### 4.2.3. Shannon entropy of receptors

The SE values for the PRAME and BAP1 sequences were clustered around 4.075*±*0.003 and 4.095*±*0.028, respectively (Table 6). As the SE values approach the theoretical maximum of 4.32, it becomes evident that a notable degree of disorderliness exists in the distribution of amino acid frequencies within both PRAME and BAP1 sequences [28]. This high level of disorderliness may bear implications for the structural and functional properties of these proteins [28, 46].

**Table 6:** Shannon entropy among the PRAME and BAP1 protein sequences.

| BAP1 | SE | PRAME | SE | PRAME | SE | PRAME | SE |
| --- | --- | --- | --- | --- | --- | --- | --- |
| BAP1_1 | 4.056 | PRAME_14 | 4.071 | PRAME_18 | 4.075 | PRAME_6 | 4.081 |
| BAP1_4 | 4.078 | PRAME_10 | 4.071 | PRAME_15 | 4.075 | PRAME_8 | 4.081 |
| BAP1_2 | 4.082 | PRAME_16 | 4.072 | PRAME_17 | 4.075 | PRAME_7 | 4.081 |
| BAP1_8 | 4.084 | PRAME_3 | 4.073 | PRAME_20 | 4.075 |  |  |
| BAP1_6 | 4.088 | PRAME_12 | 4.073 | PRAME_1 | 4.075 |  |  |
| BAP1_3 | 4.096 | PRAME_11 | 4.073 | PRAME_2 | 4.076 |  |  |
| BAP1_5 | 4.098 | PRAME_13 | 4.074 | PRAME_4 | 4.077 |  |  |
| BAP1_7 | 4.121 | PRAME_9 | 4.074 | PRAME_19 | 4.078 |  |  |
| BAP1_9 | 4.153 | PRAME_21 | 4.075 | PRAME_5 | 4.080 |  |  |

### 4.3. Homogeneous poly-string frequency of amino acids

Taking into account all amino acids across all PRAME sequences, the possible lengths of homogeneous poly-strings found out to be 1, 2 and 3. In a similar way, in BAP1 sequences the possible lengths found out to be 1, 2, 3, 4, and 6 (Table 7). In each sequence, the counts of homogeneous poly-strings for each of the twenty amino acids separately are provided in **Supplementary file 1**. It was noticed that for both types of sequences, poly-strings of length 1 were present for all twenty amino acids (**Supplementary 1**).

**Table 7:** Frequency of homogeneous poly-string of length 1, 2, … 10 for each protein sequence.

| PRAME Homogenous polystring of length 1 |  |  |  | PRAME Homogenous polystring of length 1 |  |  |  | BAP1 Homogenous polystring of length 1 |  |  |  |  |  |
| --- | --- | --- | --- | --- | --- | --- | --- | --- | --- | --- | --- | --- | --- |
| PRAME | l=1 | l=2 | l=3 | PRAME | l=1 | l=2 | l=3 | BAP1 | l=1 | l=2 | l=3 | l=4 | l=6 |
| PRAME_1 | 452 | 25 | 3 | PRAME_12 | 449 | 24 | 4 | BAP1_1 | 531 | 34 | 3 | 1 | 1 |
| PRAME_2 | 452 | 25 | 3 | PRAME_13 | 449 | 24 | 4 | BAP1_2 | 582 | 39 | 5 | 1 | 1 |
| PRAME_3 | 452 | 25 | 3 | PRAME_14 | 449 | 24 | 4 | BAP1_3 | 624 | 46 | 4 | 0 | 0 |
| PRAME_4 | 452 | 25 | 3 | PRAME_15 | 449 | 24 | 4 | BAP1_4 | 612 | 42 | 5 | 0 | 0 |
| PRAME_5 | 453 | 26 | 2 | PRAME_16 | 456 | 25 | 1 | BAP1_5 | 623 | 47 | 4 | 0 | 0 |
| PRAME_6 | 451 | 26 | 2 | PRAME_17 | 456 | 25 | 1 | BAP1_6 | 617 | 46 | 6 | 0 | 0 |
| PRAME_7 | 451 | 26 | 2 | PRAME_18 | 453 | 25 | 2 | BAP1_7 | 588 | 45 | 6 | 1 | 0 |
| PRAME_8 | 454 | 24 | 2 | PRAME_19 | 449 | 24 | 3 | BAP1_8 | 621 | 52 | 6 | 3 | 0 |
| PRAME_9 | 447 | 25 | 4 | PRAME_20 | 443 | 27 | 2 | BAP1_9 | 588 | 44 | 6 | 0 | 0 |
| PRAME_10 | 451 | 26 | 2 | PRAME_21 | 380 | 20 | 1 |  |  |  |  |  |  |
| PRAME_11 | 446 | 27 | 3 |  |  |  |  |  |  |  |  |  |  |

No PRAME sequence exhibits homogeneous poly-strings of length two for the following six amino acids: glycine, isoleucine, methionine, asparagine, tryptophan and tyrosine. All sequences possessed single occurrences of CC, DD and FF (poly-strings of length two for cysteine, aspartic acid and phenylalanine respectively). All PRAME sequences contained AA (poly-strings of alanine having length two) with count 3 except PRAME_21 which had single occurrence of AA. Single occurrence of PP (poly-strings of proline having length two) appeared in all sequences except PRAME_9 and PRAME_21 in which 2 and no PP exist respectively. All sequences composed of single occurrence of TT (poly-strings of threonine having length two) except PRAME_8, PRAME_16 and PRAME_17 in which no TT exist. Only 4 out of 21 sequences contained VV (poly-strings of valine having length two) with count one and no other sequence had any VV.

Homogeneous poly-strings of length three were absent in all PRAME sequences for fifteen amino acids. The maximum count of poly-strings of length three for each of the remaining five amino acids (glutamic acid, lysine, leucine, arginine, and serine) was 1 in any PRAME sequence and each PRAME sequence contains at least one poly-string of length three. EEE, RRR and SSS (poly-strings having length three of glutamic acid, arginine and serine respectively), were identified in 10, 11 and 13 out of 21 PRAME sequences respectively. KKK (poly-strings of lysine having length three) was found only in PRAME_11 and PRAME_20. Each PRAME sequence contained a single occurrence of LLL (poly-strings of leucine having length three) except PRAME_16 and PRAME_17 which did not exhibit any LLL. PRAME_16, PRAME_17 and PRAME_21 had only one poly-string of length three which in PRAME_16 and PRAME_17 corresponds to RRR and in PRAME_21, corresponds to LLL. Seven PRAME sequences displayed two poly-strings of length three and none of them contained EEE. Excluding PRAME_11, rest five of the six PRAME sequences, which had three poly-strings of length three, had single occurrences of EEE, LLL and SSS. Apart from LLL, other two poly-strings were KKK and RRR in PRAME_11. Each of the five PRAME sequences, which contained four poly-strings of length 3, had single occurrences of EEE, LLL, RRR and SSS. In all PRAME sequences either EE or EEE exist. Except for 3 sequences, each sequence had nine numbers of LL and one LLL. The count of LL in PRAME_16 and PRAME_17 was ten while in PRAME_21 it was seven.

Analysis of BAP1 sequences revealed only BAP1_1 and BAP1_2 had one poly-string of length six and it corresponds to glutamic acid. Single occurrences of poly-strings of length four in BAP1_1 and BAP1_ 2 correspond to PPPP while the same in BAP1_7 is of EEEE. BAP1_8 comprises of EEEE, KKKK and SSSS each with count one. No poly-string of length three was identified for the following twelve amino acids in any BAP1 sequence (C, F, H, I, K, L, M, N, Q, V, W and Y). Contribution of each amino acid reveals that only EEE had count 3 in BAP1_9 while maximum count of poly-strings of length three for all other seven amino acids was two in any sequence. Both AAA and GGG were present only in BAP1_7 and BAP1_9, each with count one. No poly-string of length two exists for following four amino acids in any BAP1 sequence (C, M, W, and Y). All BAP1 sequences had a single FF. Maximum count of poly-strings of length two for any amino acid was nine which was observed in BAP1_6 for serine.

### 4.4. Polar, non-polar residue profiles of moonlighting proteins

The percentage distributions of polar and non-polar residues were calculated for each PRAME (Figure 9, Left) and BAP1 (Figure 9, Right) sequence. The analysis revealed that the average ratio of polar to non-polar residues in PRAME sequences was 0.85 *±* 0.01, ranging from 0.83 to 0.89. In contrast, the average ratio of polar to non-polar residues in BAP1 sequences was 1.13 *±* 0.05, with a range of 1.07 to 1.2. The distinctive ratio of polar and non-polar residues in PRAME and BAP1 sequences portray their individual signature characteristics.

**Figure 9:**
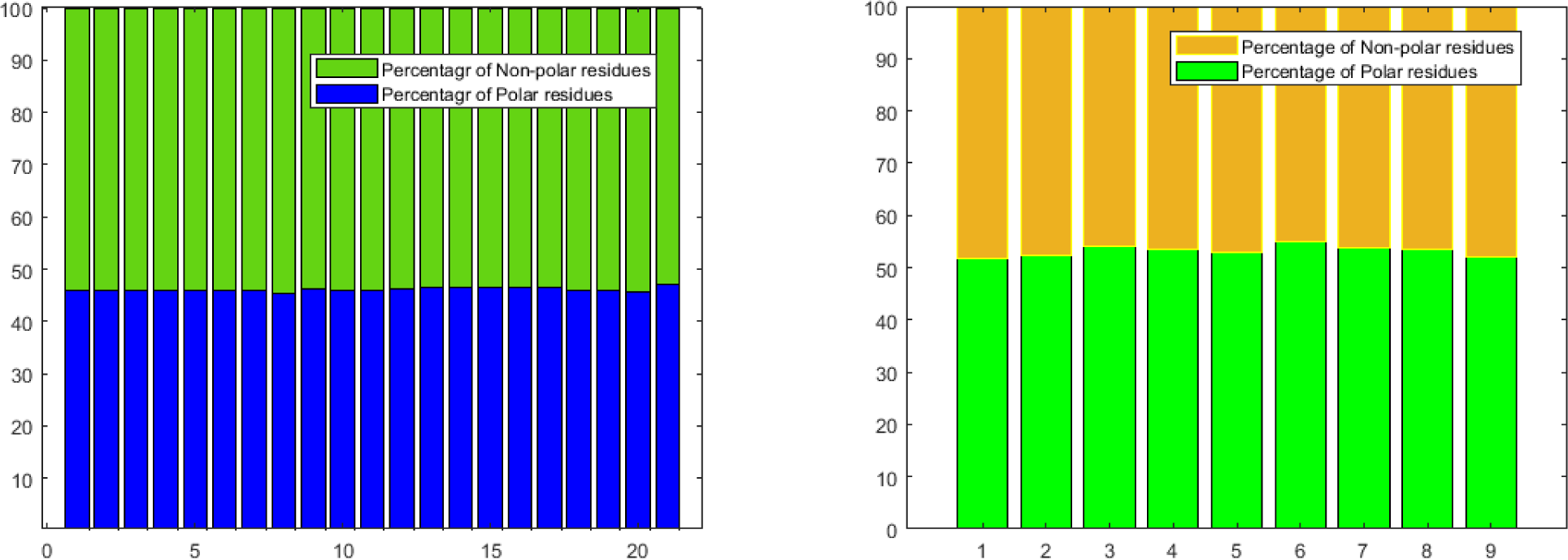
Percentages of polar, non-polar residues in PRAME (Left) and BAP1 (Right) proteins.

#### 4.4.1. Change response sequences of polar, non-polar profiles in PRAME/BAP1 proteins

Figure 10 (Left) displays the relative frequency distribution of ‘PN’, ‘NP’, ‘PP’, and ‘NN’ within the PRAME sequences. Noteworthy is the significantly high occurrence of ‘NN’ (ranging from 23.46 to 26.18) compared to ‘PP’ (16.37 to 18.11) (Figure 10 (Left)) which is due to the ratio of non-polar to polar residue ranging between 1.15 to 1.2. In case of BAP1 sequences, Figure 10 (Right) reveals notably high percentages of ‘PP’, ranging from 27.41% to 31.54%.

**Figure 10:**
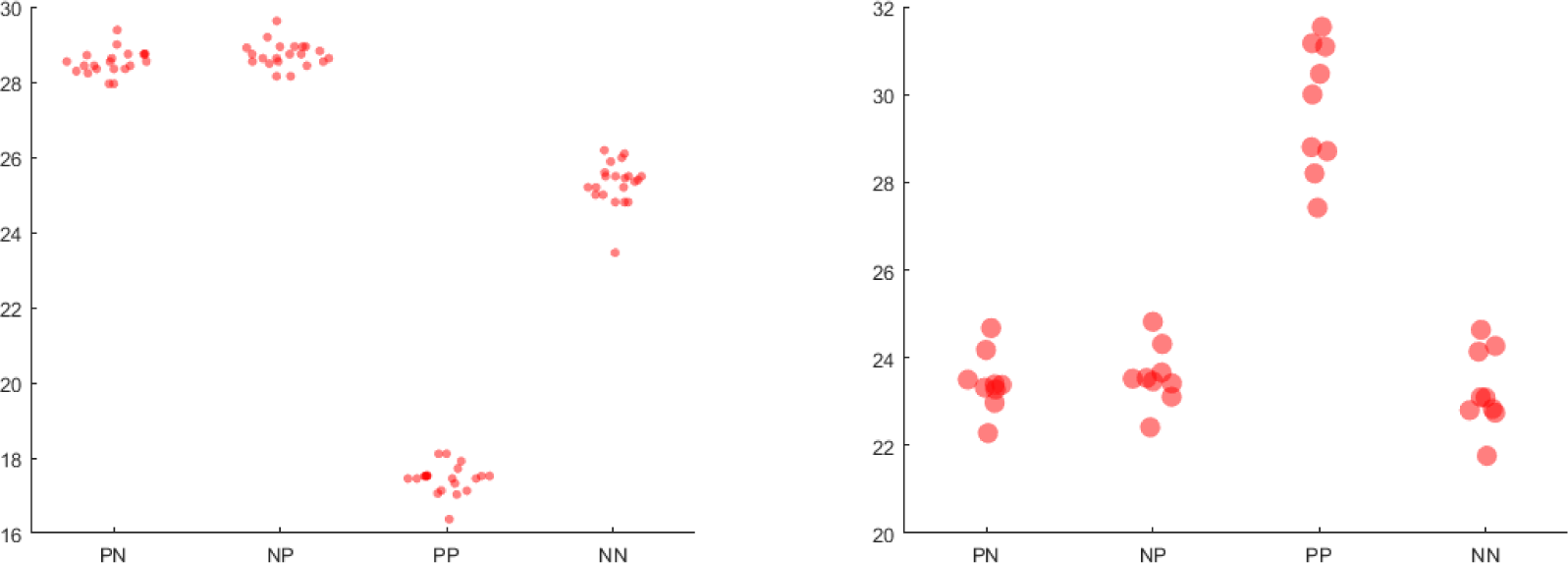
Swarm-plot of the relative frequency of PN, NP, PP, and NN changes in PRAME (left) and BAP1 (right) proteins.

Phylogenetic relationships were established by analyzing change response percentages in PRAME (Figure 11 (Top)) and BAP1 (Figure 11 (Bottom)) sequences. Notably, identical change responses were observed in three sets of proximal sets ({2, 3, 4}, {9, 12}, {13, 14, 15}, and {16, 17}), as indicated in Scale-0 (Table 8). PRAME_21 was distant from the rest of the sequences. Additionally, BAP1_8 was not included in any proximal set till the threshold we considered, as outlined in Table 8.

**Figure 11:**
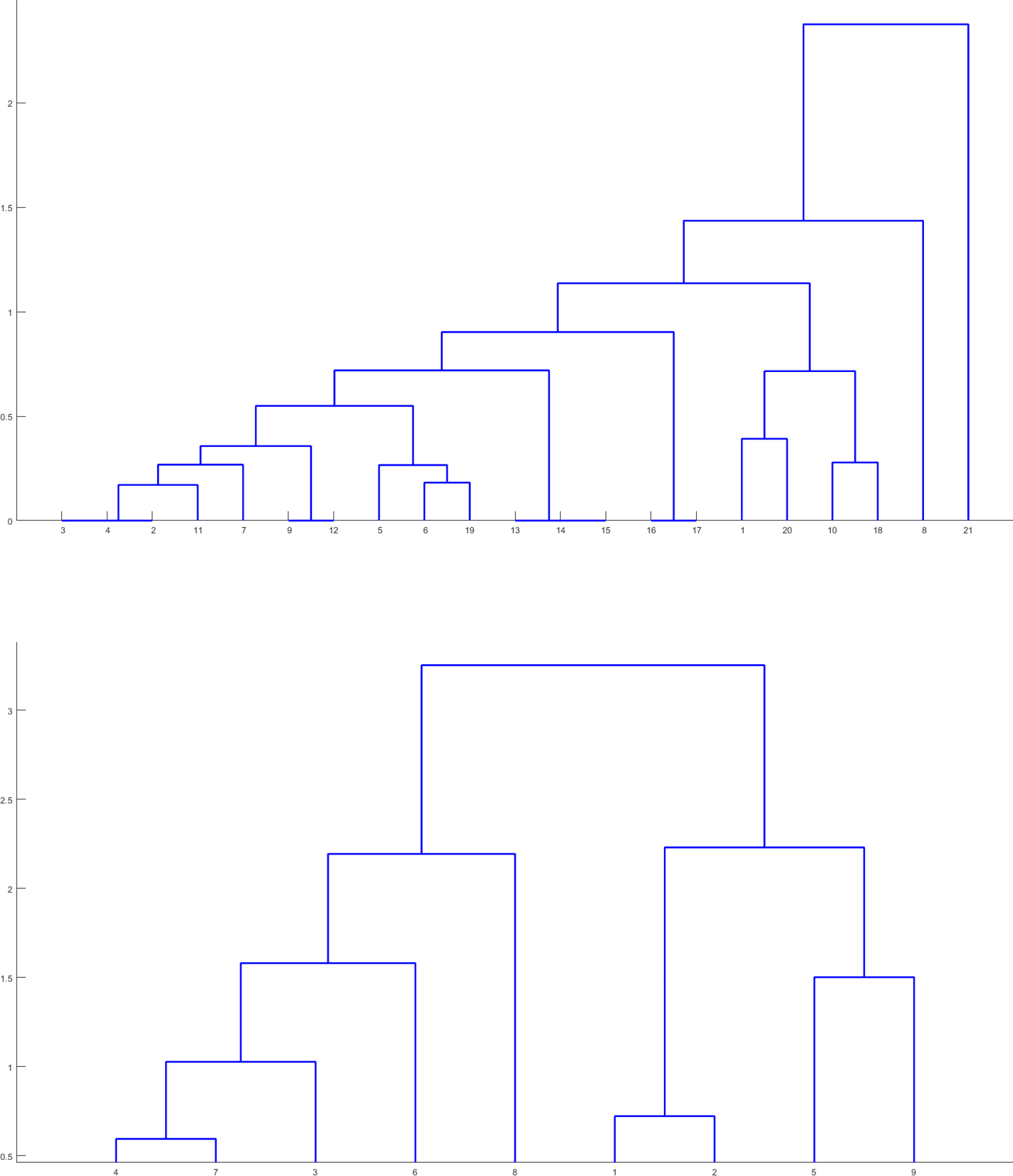
Phylogenetic relationship among the PRAME (Top) and BAP1 (Bottom) proteins based on relative frequency of PP, NP, PP, and NN changes as obtained from polar, non-polar profiles.

**Table 8:**
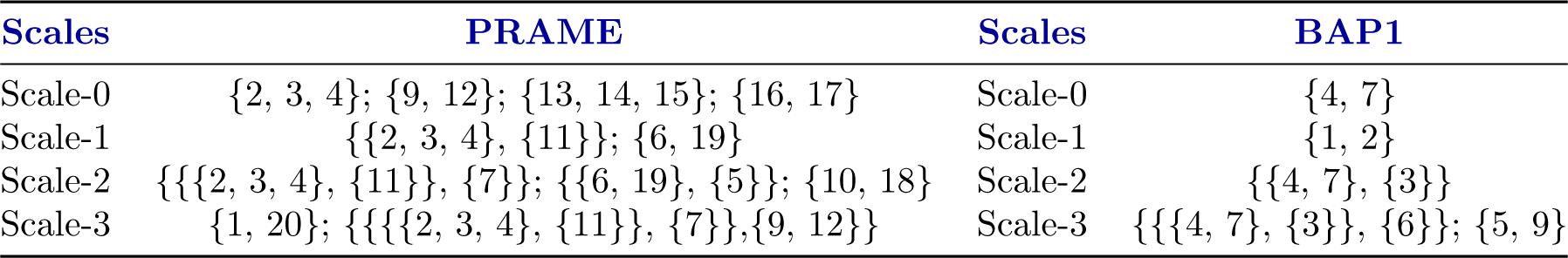
Scaled-based proximal sets of PRAME/BAP1 proteins based on change response of polar and non-polar residues compositions.

### 4.5. Acidic, basic, neutral residue based phylogenetic relationship

Figure 12 revealed percentages of neutral residues ranging from 78.58% to 80.03% for PRAME and from 70.59% to 74.49% for BAP1. The ratio of acidic to basic residue of the PRAME sequences was 1.11*±* 0.05, while for BAP1 sequences, it was 1.04 *±* 0.06. This suggests a relatively balanced distribution between acidic and basic residues across PRAME and BAP1 sequences.

**Figure 12:**
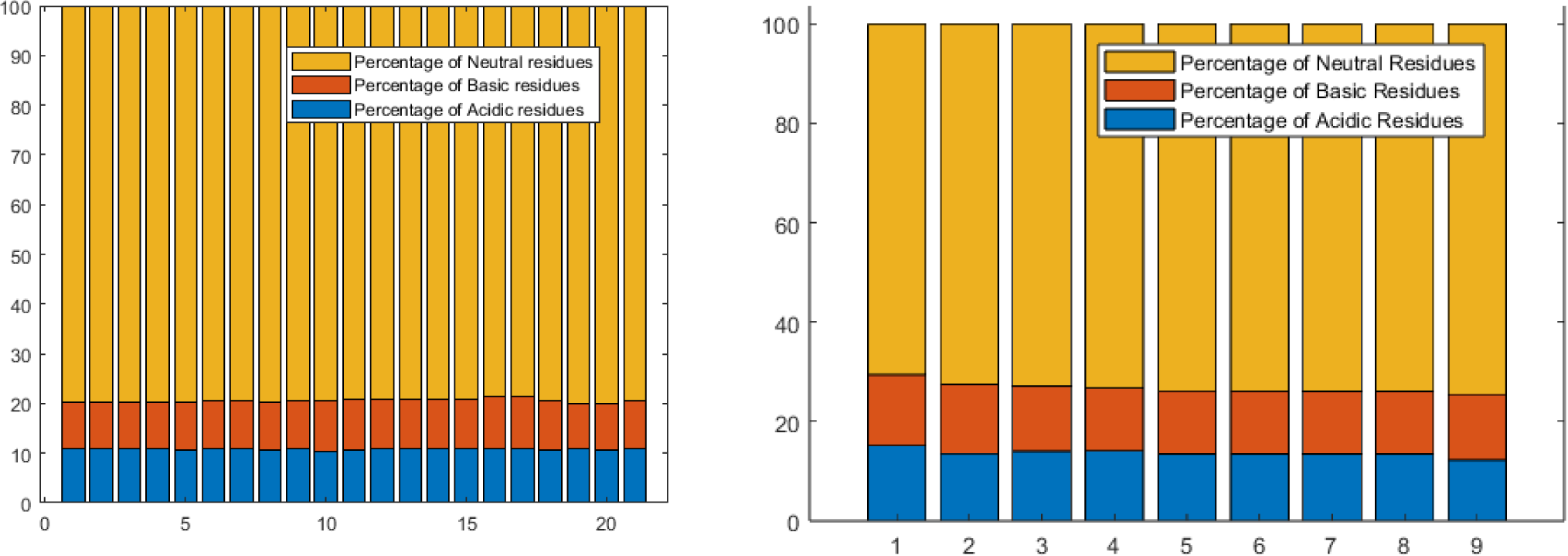
Percentages of acidic, basic, neutral residues in PRAME (Top) and BAP1 (Bottom) proteins.

#### 4.5.1. Change response sequences of acidic, basic, and neutral profiles

As percentages of neutral residues were significantly higher compared to acidic or basic, the percentage of ‘NN’ in all PRAME sequences was significantly higher (ranging from 62.6% to 65.35%) than other eight changes (Figure 13). The ratios of ‘NA’ and ‘AN’ lay between 0.98 and 1.06 while ratios of ‘NB’ and ‘BN’ varied from 0.93 to 1.03. Hence, an almost similar number of changes from acidic (basic) to neutral and neutral to acidic (basic) residues was noted in all PRAME sequences. Ratios of ‘NA’ and ‘NB’ were turned out to be between 1.2 to 1.46 suggesting consecutive presence of basic residues was more than consecutive presence of acidic residues.

**Figure 13:**
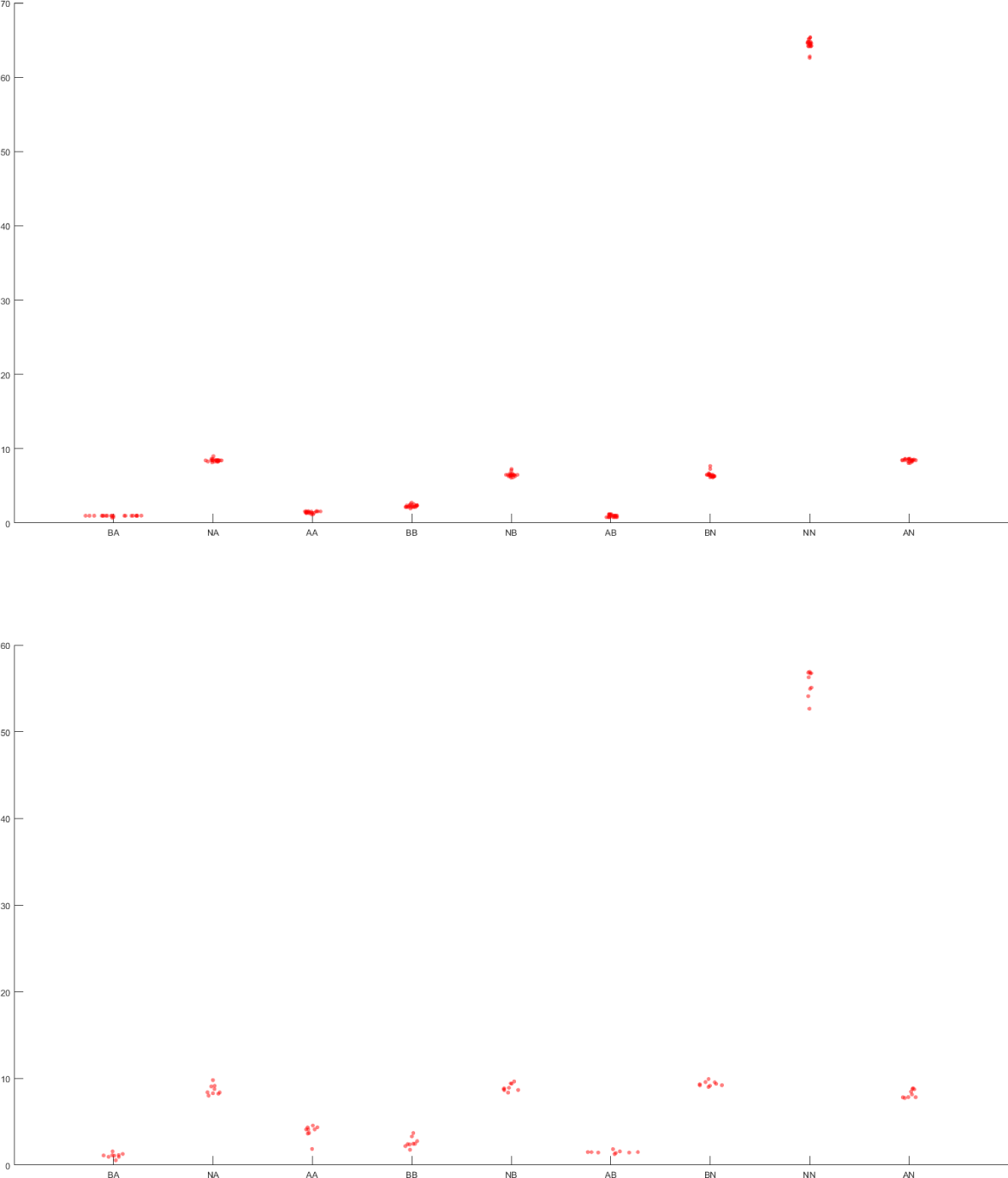
Swarm-plot of the relative frequency of all nine changes in PRAME (top) and BAP1 (bottom) proteins.

The percentage of ‘NN’ in BAP1 sequences spanned between 52.65 (in BAP1_8) and 56.9. Almost similar numbers of changes from acidic to neutral and neutral to acidic residues were noted in all BAP1 sequences. Ratio of ‘NB’ and ‘BN’ was lowest in BAP1_9 (0.89), while it varied between 0.94 to 0.98 for the other eight sequences. The ratio of ‘AA’ and ‘BB’ was the highest for BAP1_1 (2.09) as percentage of ‘BB’ was significantly lower in BAP1_1 while the ratio of ‘AA’ and ‘BB’ was lowest for BAP1_9 (0.56) as percentage of ‘AA’ was significantly lower in BAP1_9. The percentage of ‘NA’ was the highest in BAP1_9 (9.8) while the same varied between 8.03 and 9.15 for the other eight BAP1 sequences. The percentage of ‘BA’ was significantly lower in BAP1_9 compared for the other BAP1 sequences.

PRAME_1, PRAME_2, PRAME_3, and PRAME_4 had identical feature vectors consisting of relative frequency of nine changes. The same was the case for PRAME_12, PRAME_13, PRAME_14 and PRAME_15 (Figure 14 (Top)). As indicated in Table 9 PRAME_16 and PRAME_17 were proximal, but distant from the rest of the PRAME sequences.

**Figure 14:**
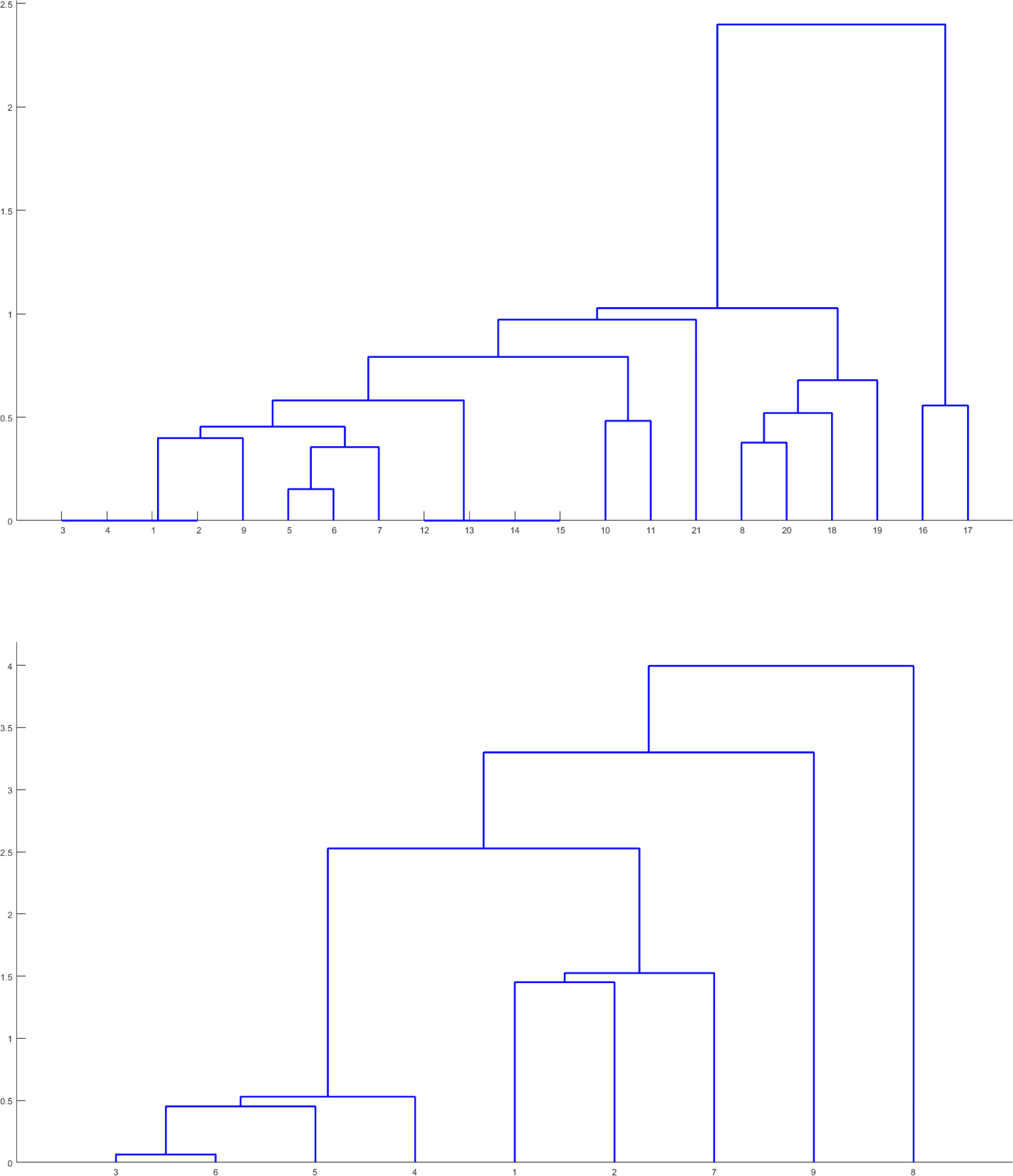
Phylogenetic relationship among the PRAME (Top) and BAP1 (Bottom) proteins based on relative frequency of BA, NA, AA, BB, NB, AB, BN, NN, and AN changes as obtained from acidic, basic, and neutral profiles.

**Table 9:** Scaled based proximal sets of PRAME/BAP1 proteins based on change response of acidic, basic, and neutral residues compositions.

| Scales | PRAME | Scales | BAP1 |
| --- | --- | --- | --- |
| Scale-0 | {1, 2, 3, 4}; {12, 13, 14, 15} | Scale-0 | {3, 6} |
| Scale-1 | {{1, 2, 3, 4}, {9}}; {{5, 6}, {7}}; {8, 20} | Scale-1 | {{3, 6}, {5}} |
| Scale-2 | {{8, 20}, {18}}; {10, 11}; {{{1, 2, 3, 4}, {9}}, {{5, 6}, {7}}} | Scale-2 | {{{3, 6}, {5}}, {4}} |
| Scale-3 | {16, 17}; {{{{1, 2, 3, 4}, {9}}, {{5, 6}, {7}}}, {12, 13, 14, 15}} | Scale-3 | {{1, 2}, {7}} |

Two BAP1 sequences (BAP1_8 and BAP1_9) were relatively distant from each other as well as mutually (Table 9). BAP1_3 and BAP1_6 were very much proximal (Figure 14 (Bottom)).

### 4.6. Intrinsic protein disorder analysis of PRAME and BAP1 sequences

Within the PRAME sequences (see Figure 15 - Top), a notable prevalence of moderately flexible residues was evident (ranging from 39.49% to 47.64%). The proportion of highly flexible residues within each PRAME sequence was centered around 35.37%, with a standard deviation of 1.97. The percentage of disordered residues was significantly lower in PRAME_7 (11.59%) than in the other PRAME sequences while the highly flexible residue was the highest in PRAME_7 (40.47%).

**Figure 15:**
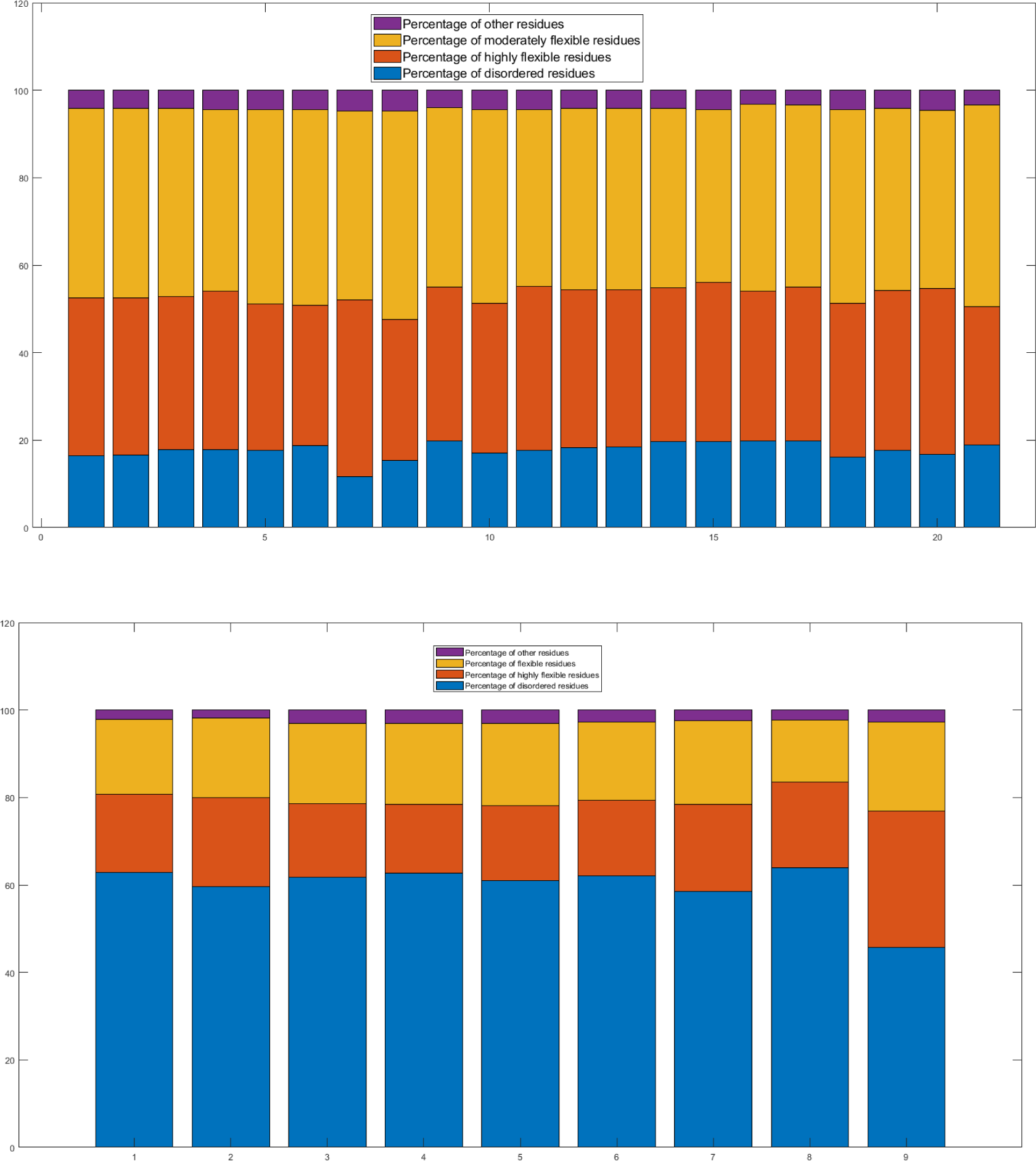
Percentages of disordered, highly flexible, moderately flexible, and other residues in PRAME (top) and BAP1 (bottom) proteins.

In the BAP1 sequences (see Figure 15 - Bottom), a distinctive region emerged as the disordered region, indicated by disordered residues ranging from 45.68% to 63.97%. BAP1_9 possessed a relatively higher percentage of highly flexible residues (31.2%) and lower percentage of disordered residues (45.68%) compared to the rest eight BAP1 sequences. The percentage of highly flexible residues for the rest eight sequences varied from 15.75 to 20.44. Among all BAP1 sequences, BAP1_8 contained the lowest percentage of moderately flexible residues (14.17%).

#### 4.6.1. Percentage of four different IPD residue types in each amino acid

Each of the four figures (Figures 16, 17, 18, and 19) corresponds to one residue type and displays the percentage of that residue in each amino acid. It needs to be pointed out that percentages were calculated based on individual amino acid frequency in a sequence implying the sum of the percentages of four different IPD residue types will be 100 for a given amino acid in a sequence.

**Figure 16:**
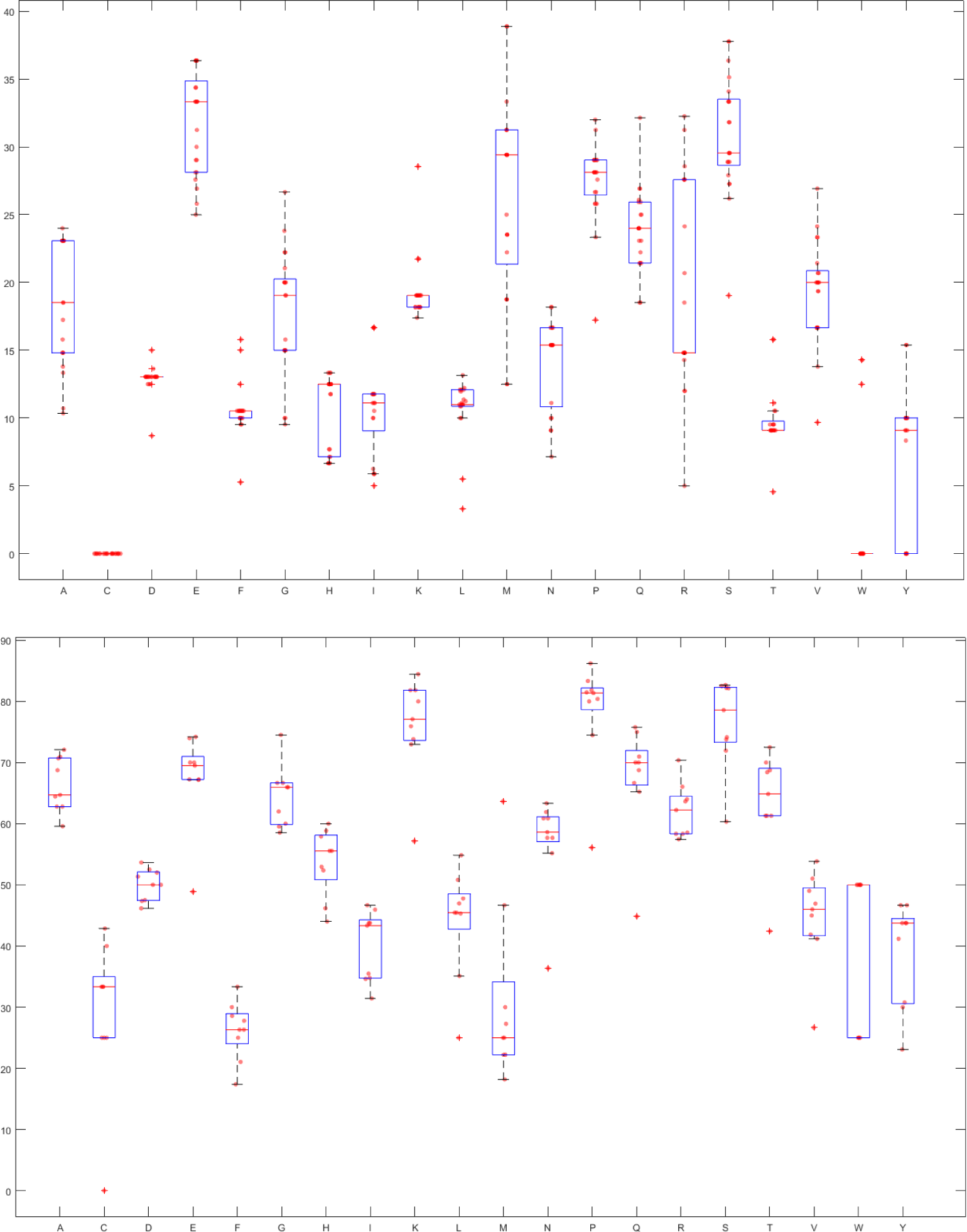
Percentage of disordered regions in each amino acid for PRAME (Top) and BAP1 (Bottom)

Figure 16 (Top) revealed that among all amino acids, glutamic acid (E) had the highest median percentage of ‘D’ residues (33.33%) in PRAME. Methionine (M), proline (P) and serine (S) had also relatively high median percentage of ‘D’ residues compared to other amino acids. Cysteine (C) had no ‘D’ residues in any sequence and Tryptophan (W) had no ‘D’ residues except three sequences (PRAME_16, PRAME_17 and PRAME_18). Across all amino acids and all sequences, maximum percentage of ‘D’ residues (38.89%) was observed for methionine followed by serine (37.78%) in both PRAME_16 and PRAME_17. Studying individual amino acids showed lowest percentage of ‘D’ residues for six amino acids (D, F, P, S, T, and V) was found in PRAME_7 among all sequences. In addition, PRAME_16 and PRAME_17 had the highest percentage of ‘D’ residues for both T and Y while the highest percentage of ‘D’ residues for both G and K was observed in PRAME_21.

Comparing different amino acids from Figure 16 (Bottom) it was noticed that median percentage of ‘D’ residues was the highest (81.36%) for proline (P) in BAP1 sequences followed by serine and lysine (K) with 78.57% and 77.08%, respectively. Methionine and phenylalanine (F) had the lowest median ‘D’ residues of 25% and 26.31%, respectively. BAP1_9 displayed some unique characteristics like no ‘D’ residues in cysteine (C) coupled with significantly low percentage of ‘D’ residues compared to other sequences.for following amino acids: alanine (A), glutamic acid (E), phenylalanine, histidine (H), lysine, leucine (L), asparagine (N), proline, glutamine (Q), serine, threonine (T), valine (V) and tyrosine (Y). Further, the percentage of ‘D’ residues in methionine was maximum in BAP1_9 among all sequences.

It was noted from Figure 17 (Top) that in PRAME sequences, aspartic acid (D) had the highest median percentage (56.5%) of ‘HF’ residues followed by cysteine (C) and asparagine (N) with 53.3% and 50% respectively. Among all amino acids of all sequences, asparagine had the highest percentage (72.73%) of ‘HF’ residues in PRAME_10. Highest percentage of ‘HF’ in W was found in PRAME_2 while no ‘HF’ residue of W was present in PRAME_16, PRAME_17 and PRAME_21. Further, W had the lowest median percentage. Among all PRAME sequences, PRAME_21 contained highest ‘HF’ percentage of A while the lowest ‘HF’ percentage of D, G, K, M and Q. Highest ‘HF’ percentage of E, F, G, Q and V was observed in PRAME_7. PRAME_1 and PRAME_2 had a significantly low ‘HF’ percentage of I compared to other sequences. PRAME_8 contained the lowest ‘HF’ percentage of C, F and T. PRAME_17 displayed the highest (lowest) ‘HF’ percentage of K (R). PRAME_18 and PRAME_21 showed much lower ‘HF’ percentage of V among all sequences.

**Figure 17:**
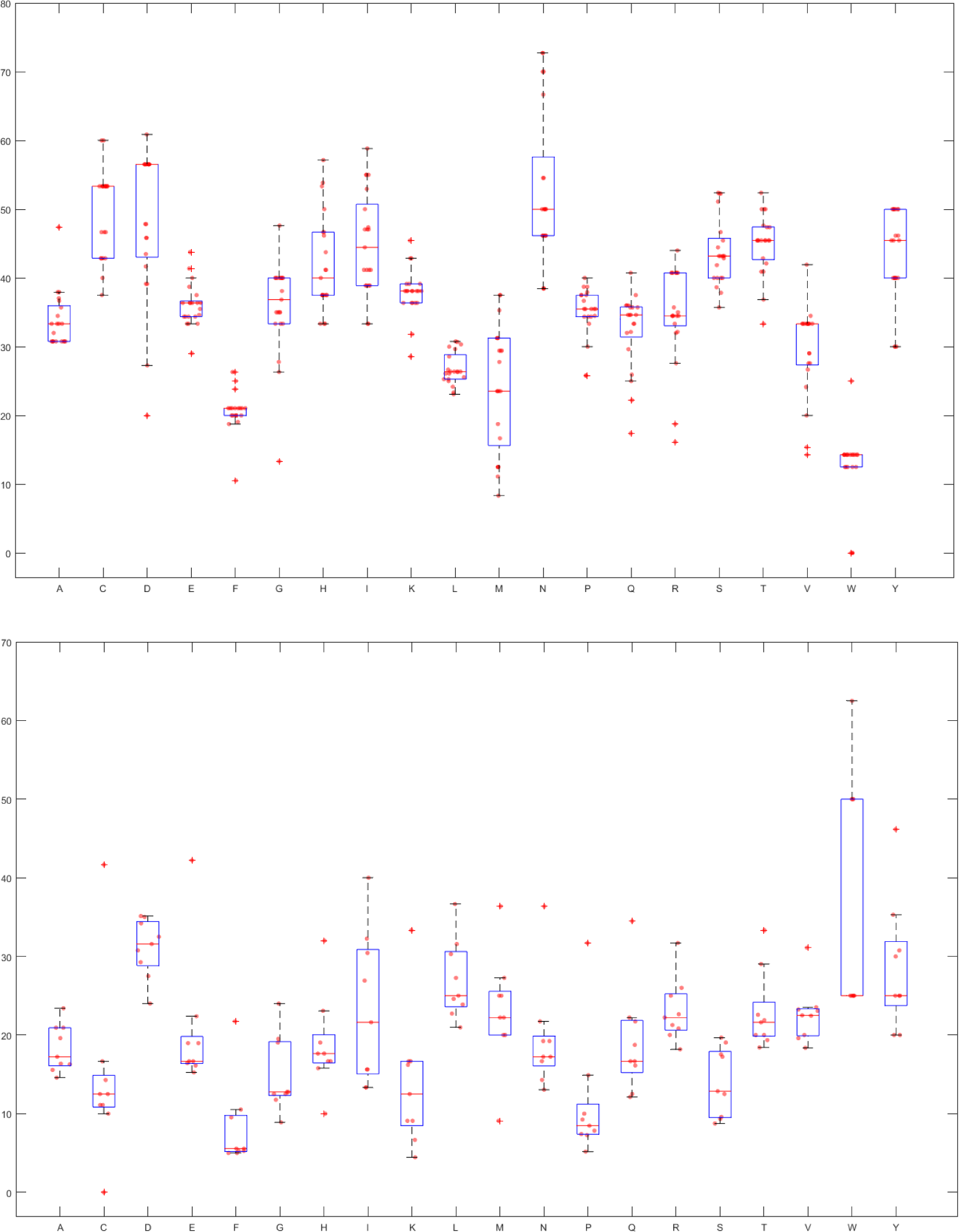
Percentage of highly flexible regions in each amino acid for PRAME (Top) and BAP1 (Bottom)

Figure 17 (Bottom) exhibited that among all amino acids, aspartic acid and phenylalanine (F) contained the highest (31.6%) and the lowest (5.55%) median percentage of ‘HF’ residues respectively in BAP1 sequences. Compared to all other BAP1 sequences, BAP1_9 had a significantly higher ‘HF’ percentage in fifteen amino acids namely cysteine, glutamic acid (E), phenylalanine (F), glycine (G), histidine (H), isoleucine (I), lysine (K), leucine (L), asparagine (N), proline (P), glutamine (Q), threonine (T), valine (V), tryptophan and tyrosine (Y) while significantly lower ‘HF’ percentage in methionine (M). BAP1_7 had no ‘HF’ residue of cysteine and a significantly higher ‘HF’ percentage in methionine than other sequences. Considering all amino acids across all sequences, a maximum percentage (62.5%) of ‘HF’ residues was observed in tryptophan for BAP1_9.

Figure 18 (Top) indicated that among all amino acids of all PRAME sequences, tryptophan (W) in PRAME_21 contained the highest percentage (80%) of ‘MF’ residues and this percentage was significantly higher than maximum ‘MF’ percentage of any other amino acid. The minimum percentage (13.33%) of ‘MF’ residues was seen for serine (S) in PRAME_16. Median percentage of ‘MF’ was the highest for Leucine (L) followed by tryptophan (W) with values 58.24% and 57.14% respectively. The lowest ‘MF’ percentage of A and the highest ‘MF’ percentage of D, G, Q and R was obtained in PRAME_21. PRAME_11 and PRAME_20 both had lowest ‘MF’ percentage of C among all sequences. Highest ‘MF’ percentage of E and V was noticed in PRAME_18. Highest and lowest ‘MF’ percentage of F was found in PRAME_16 and PRAME_15 respectively. Maximum ‘MF’ percentage of H, K, L and T was obtained in PRAME_8. Minimum ‘MF’ percentage of H, I, K was present in PRAME_17. Lowest ‘MF’ percentage of Y correspond to PRAME_16 and PRAME_17.

**Figure 18:**
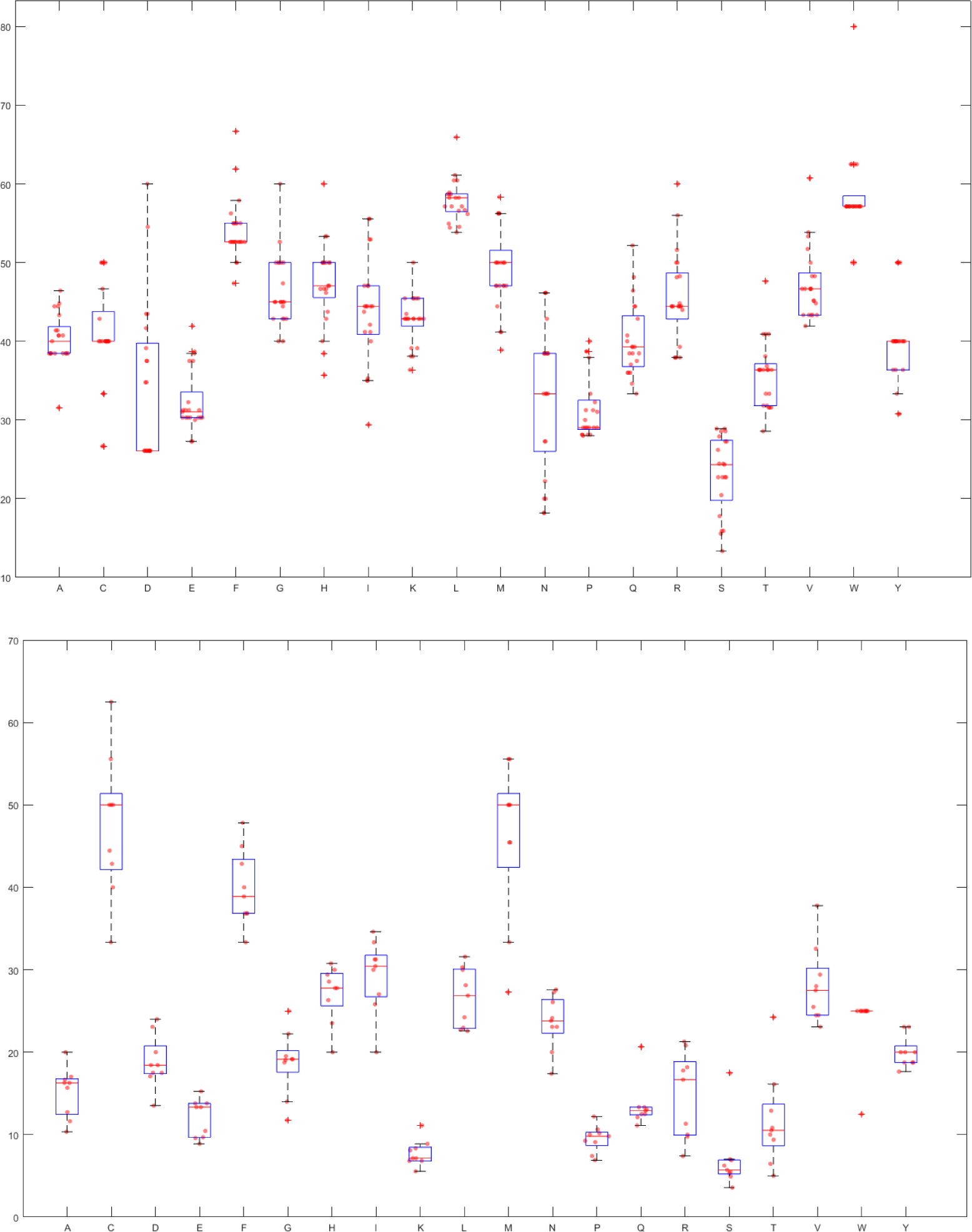
Percentage of moderately flexible regions in each amino acid for PRAME (Top) and BAP1 (Bottom)

Cysteine (C) and methionine (M) contained the highest median percentage (both 50%) of ‘MF’ residues in BAP1 sequences (Figure 18 (Bottom)). All other amino acids had significantly lower median than these two. Serine (S) had the lowest median of 5.71% followed by lysine (K) having median 7.14%. Similar to other residue types, BAP1_9 here also showed distinct characteristics like ‘MF’ percentages of phenylalanine (F), glutamine (Q), serine, threonine (T) and valine (V) were significantly higher while ‘MF’ percentages of isoleucine (I), methionine (M) and tryptophan (W) were quite lower than rest of the sequences. ‘MF’ percentage in cysteine (62.5%) of BAP1_7 was the maximum ‘MF’ percentage of all amino acids and sequences.

It was observed from Figure 19 (Top) that glutamic acid (E), histidine (H), isoleucine (I), lysine (K), asparagine (N) and arginine (R) contained no ‘O’ residue in any PRAME sequence. Tryptophan (W) had a maximum median percentage of ‘O’ (28.57%) which was significantly higher than rest amino acids. Glycine (G) had ‘O’ residue in only PRAME_9. Methionine (M) had ‘O’ residue in only PRAME_16 and PRAME_17. Further, significantly low ‘O’ percentage in phenylalanine (F) was found in PRAME_16 and PRAME_17 compared to other PRAME sequences. No ‘O’ residue of valine (V) was present in PRAME_16 and PRAME_17. Maximum ‘O’ percentage of Q was noticed in PRAME_8 while Q had no ‘O’ residue in PRAME_9, PRAME_16 and PRAME_17. Only PRAME_21 did not contain any ‘O’ residue of proline (P). Additionally, PRAME_21 had relatively low ‘O’ percentage in threonine (T) compared to other sequences. Figure 19 (Bottom) shows that none of the BAP1 sequences contained ‘O’ residue of aspartic acid (D), glutamic acid (E), methionine (M), asparagine (N), glutamine (Q), arginine (R) and tryptophan (W). Non zero ‘O’ percentage in alanine (A) and histidine (H) was only present in BAP1_8 and BAP1_9 respectively. ‘O’ residue of valine (V) and Glycine (G) was only absent in BAP1_8 and BAP1_9 respectively. Additionally, BAP1_9 and BAP1_1 had significantly higher ‘O’ percentage in leucine (L) and tyrosine (Y) compared to other sequences. Maximum median ‘O’ percentage (30%) was found in phenylalanine (F) among all amino acids.

**Figure 19:**
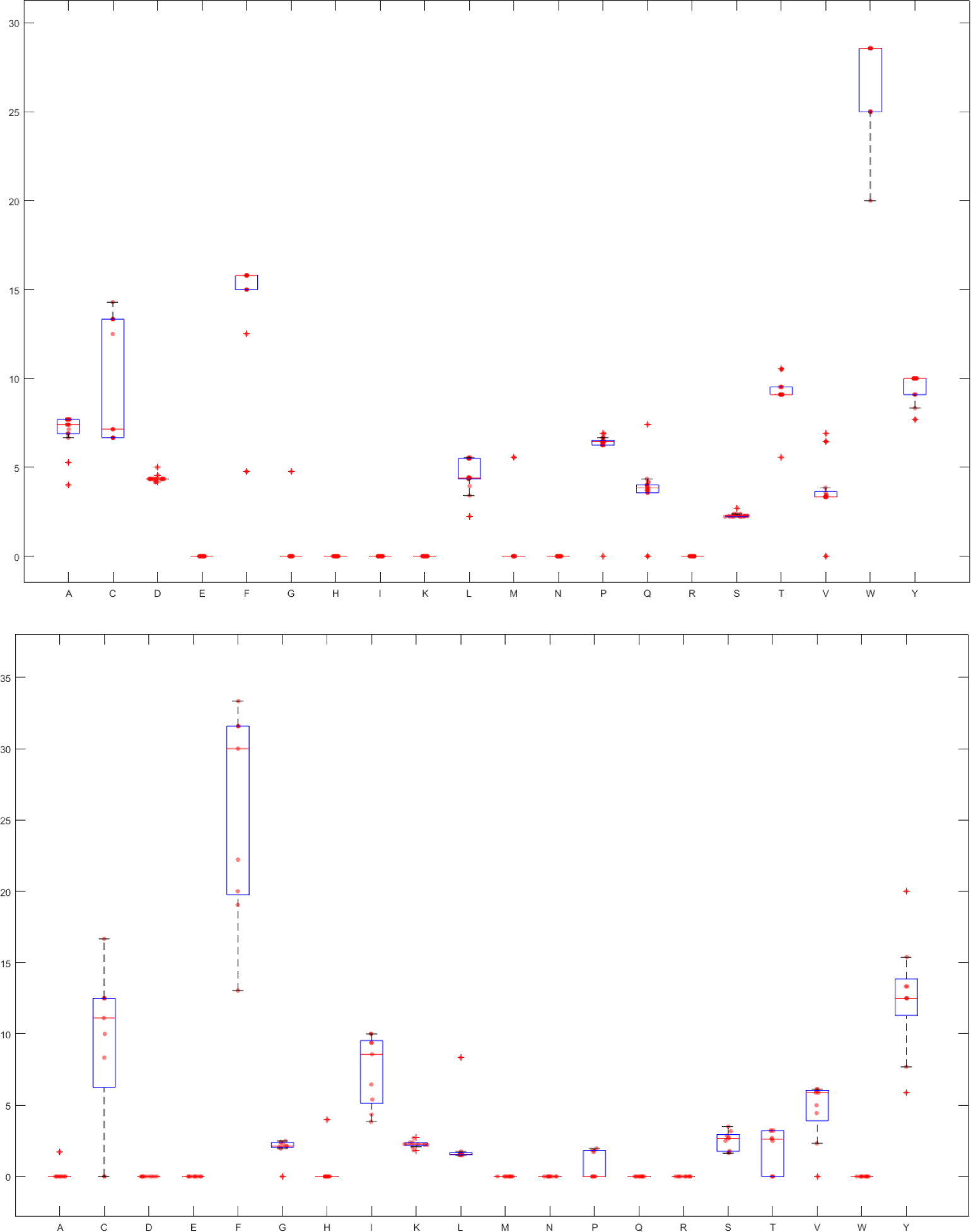
Percentage of amino acids in other regions of PRAME (Top) and BAP1 (Bottom)

#### 4.6.2. Change response sequences of disordered, highly flexible, moderately flexible, and other residues profiles

Figure 20 showed that there were no change responses for the transitions O_HF, O_D, MF_D, HF_O, D_O, and D_MF in any of the PRAME or BAP1 sequences.

**Figure 20:**
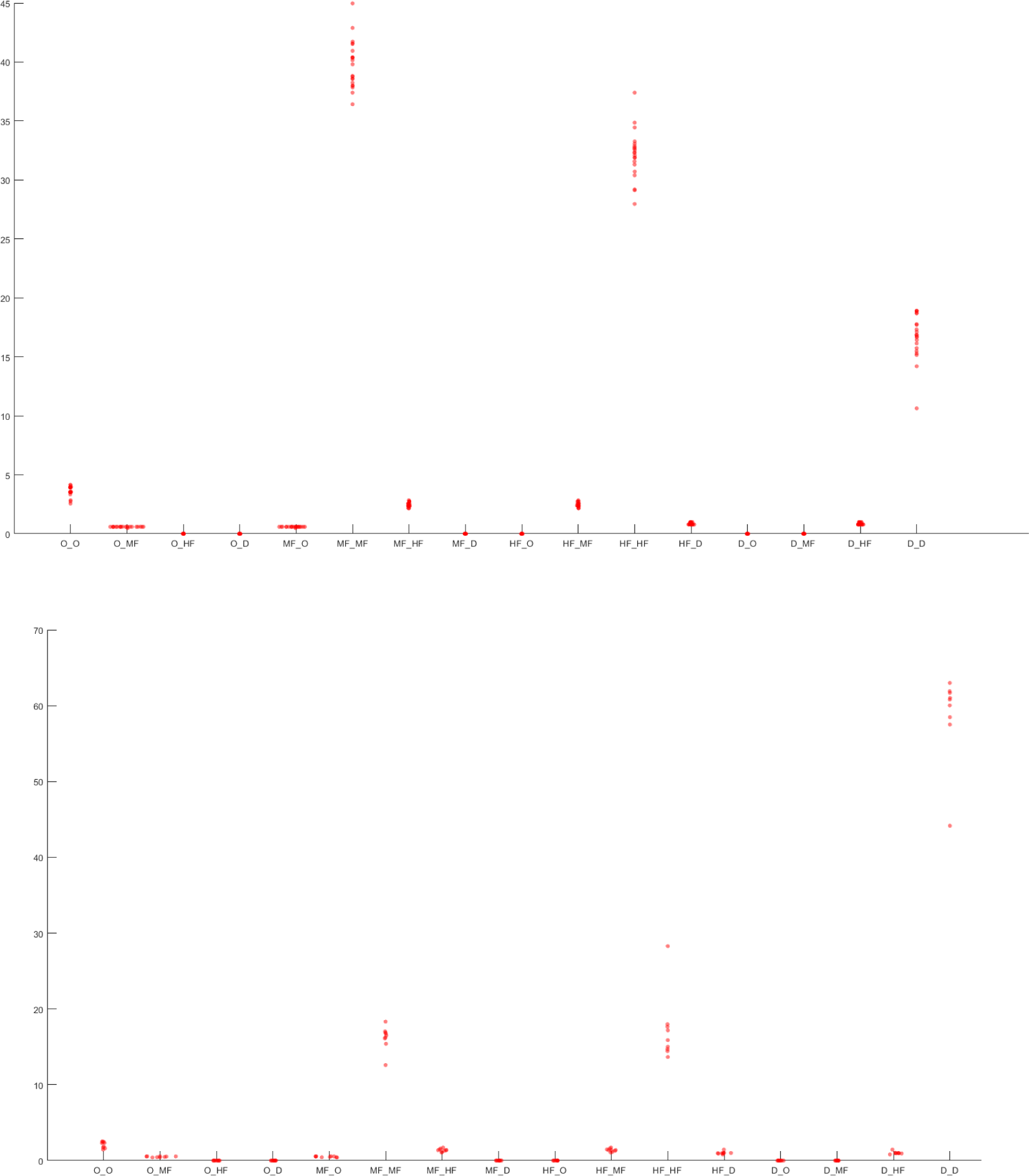
Swarm-plot of the percentages of disordered, highly flexible, moderately flexible and other residues changes in PRAME (top) and BAP1 (bottom) protein sequences.

Among all sixteen possible changes, the highest proportions were found for MF_MF in each PRAME sequence (Figure 20 (Top)). Maximum MF_MF (44.97%) and maximum HF_HF (37.4%) were noted for PRAME_8 and PRAME_7 respectively. PRAME_7 contained minimum percentage of D_D (10.63%).

Figure 20 (Bottom) displayed that median values of MF_MF and HF_HF were almost equal with values 16.5 and 15.88 respectively. BAP_9 possessed unique characteristics like much lower D_D (44.14%) and much higher HF_HF (28.28%) compared to other BAP1 sequences.

Three sequences namely PRAME_15, PRAME_8 and PRAME_7 were not included in any proximal set till the scale we considered (Figure 21 (Top) and Table 10). Notably, PRAME_7 was significantly distant from all of the other PRAME sequences. On the other hand, BAP1_9 was distant from all other eight BAP1 sequences.

**Figure 21:**
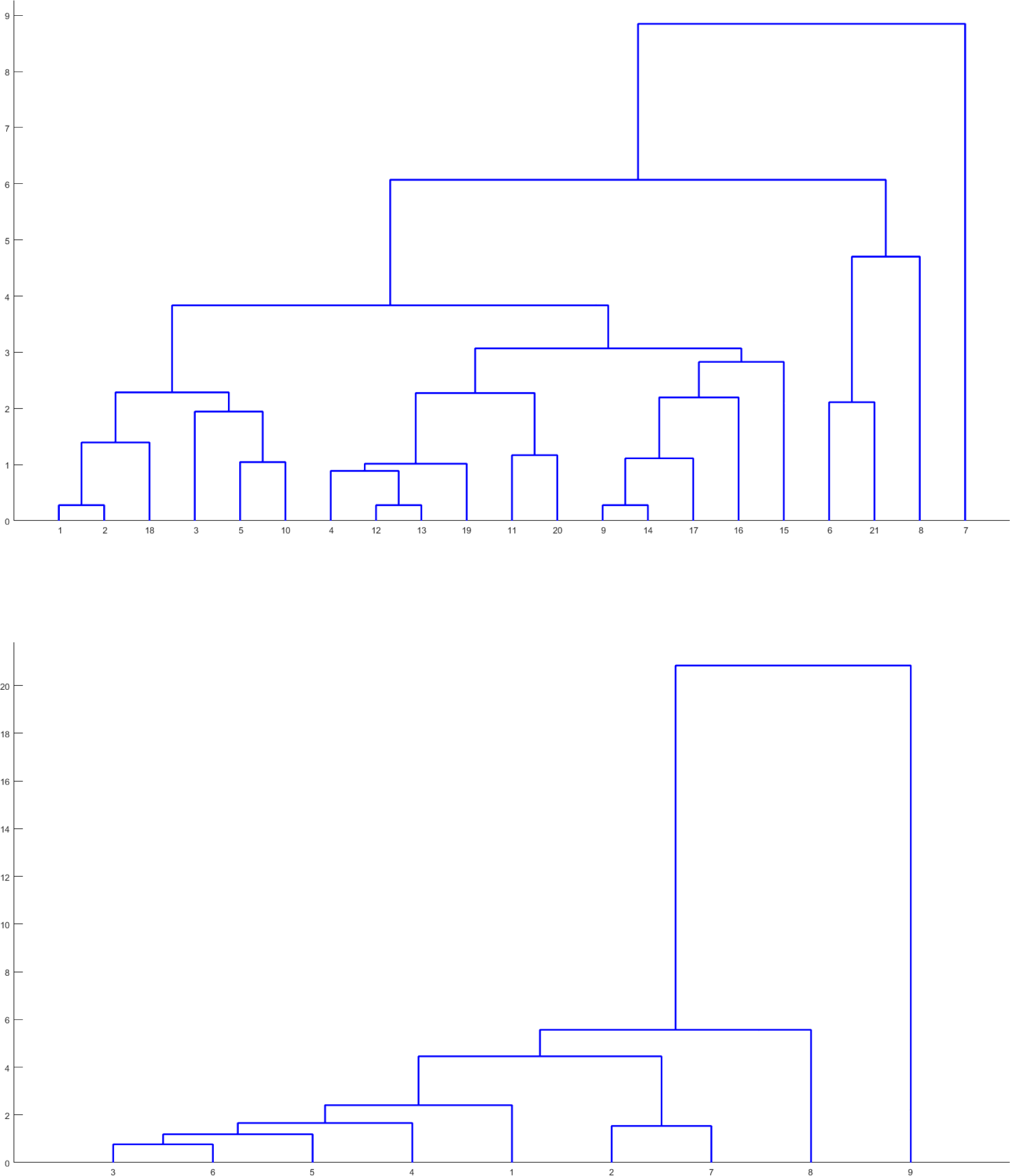
Phylogenetic relationship among the PRAME (top) and BAP1 (bottom) proteins based on relative frequency of sixteen changes as obtained from disordered, highly flexible, moderately flexible and other residues profile.

**Table 10:**
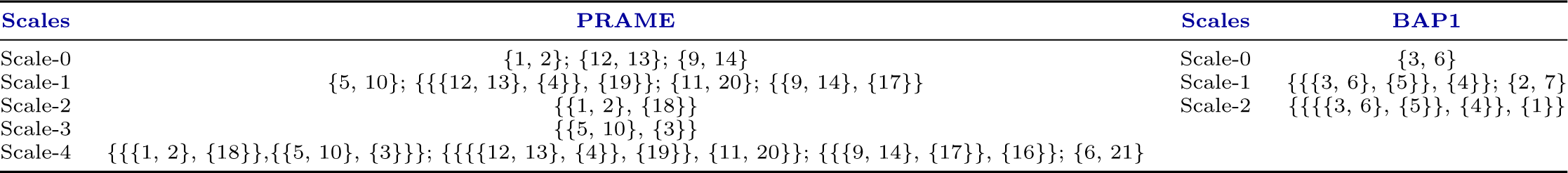
Scaled-based proximal sets of PRAME/BAP1 proteins based on change response of intrinsic protein disordered regions.

#### 4.6.3. Functional disorder analysis of human PRAME and BAP1 proteins

Functional intrinsic disorder status of human BAP1 and PRAME proteins is illustrated by Figure 22, 23. Analysis of these data indicated that proteins are located at the two poles of intrinsic disorder spectrum, with PRAME being mostly ordered (Figure 22A(Top)), and with BAP1 showing high intrinsic disorder propensity (Figure 22A(Bottom)). In fact, the PONDR® VSL2-predicted disorder contents (PDCs) of PRAME and BAP1 are 16.1% and 61.0%, and their mean disorder scores (MDSs) are 0.31 *±* 0.17 and 0.58 *±* 0.3, respectively. In other words, based on the criteria accepted in the field, where two arbitrary cutoffs for the levels of intrinsic disorder are used to classify proteins as highly ordered (*PDC <* 10%), moderately disordered (10% *≤ PDC <* 30%) and highly disordered (*PDC ≥* 30%)), human PRAME and BAP1 are clearly classified as mostly ordered (moderately disordered) and highly disordered proteins, respectively [42]. Disorder scores of significant portion (70.7%) of human PRAME residues lie between 0.15 and 0.5 i.e. ordered but flexible regions(Figure 22A(Top)). In line with these predictions, Figure 23A(left) shows that structure of most of the human PRAME1 can be predicted with high confidence, whereas Figure 23A(right) shows that structure of many regions of BAP1 can be predicted with low or very low confidence, providing further support to their disorder nature.

**Figure 22:**
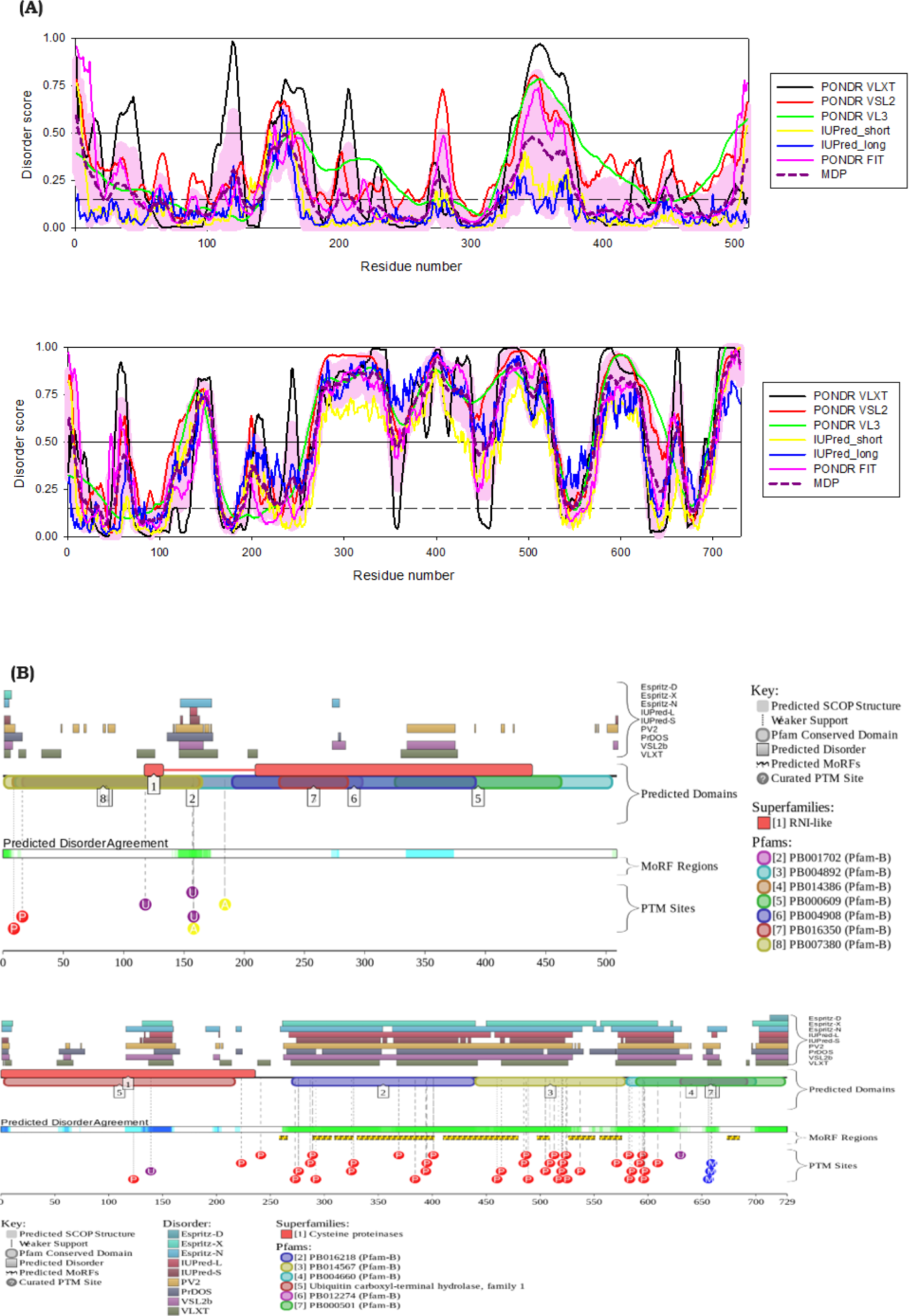
Evaluation of functional intrinsic disorder in human PRAME (UniProt ID: P78395) and human BAP1 (UniProt ID: Q92560). A. Disorder profile assembled using RIDAO that assembles outputs of six commonly used PONDR® VLS2, PONDR® VL3, PONDR® VLXT, PONDR® FIT, IUPred-Long, and IUPred-Short [41, 36, 48, 39, 40, 49, 35]. Mean disorder profile (MDP) is calculated by averaging the outputs of individual predictors is shown by dashed dark cyan curve, whereas light cyan shade represents the distribution of standard deviations. Horizontal lines represent flexibility (dashed line at disorder score 0.15) and intrinsic disorder (solid line at disorder score 0.5) thresholds. B. Functional disorder profile generated by *D*^2^*P* ^2^ [50]. In both A and B, top and bottom figures correspond to human PRAME and BAP1, respectively.

**Figure 23:**
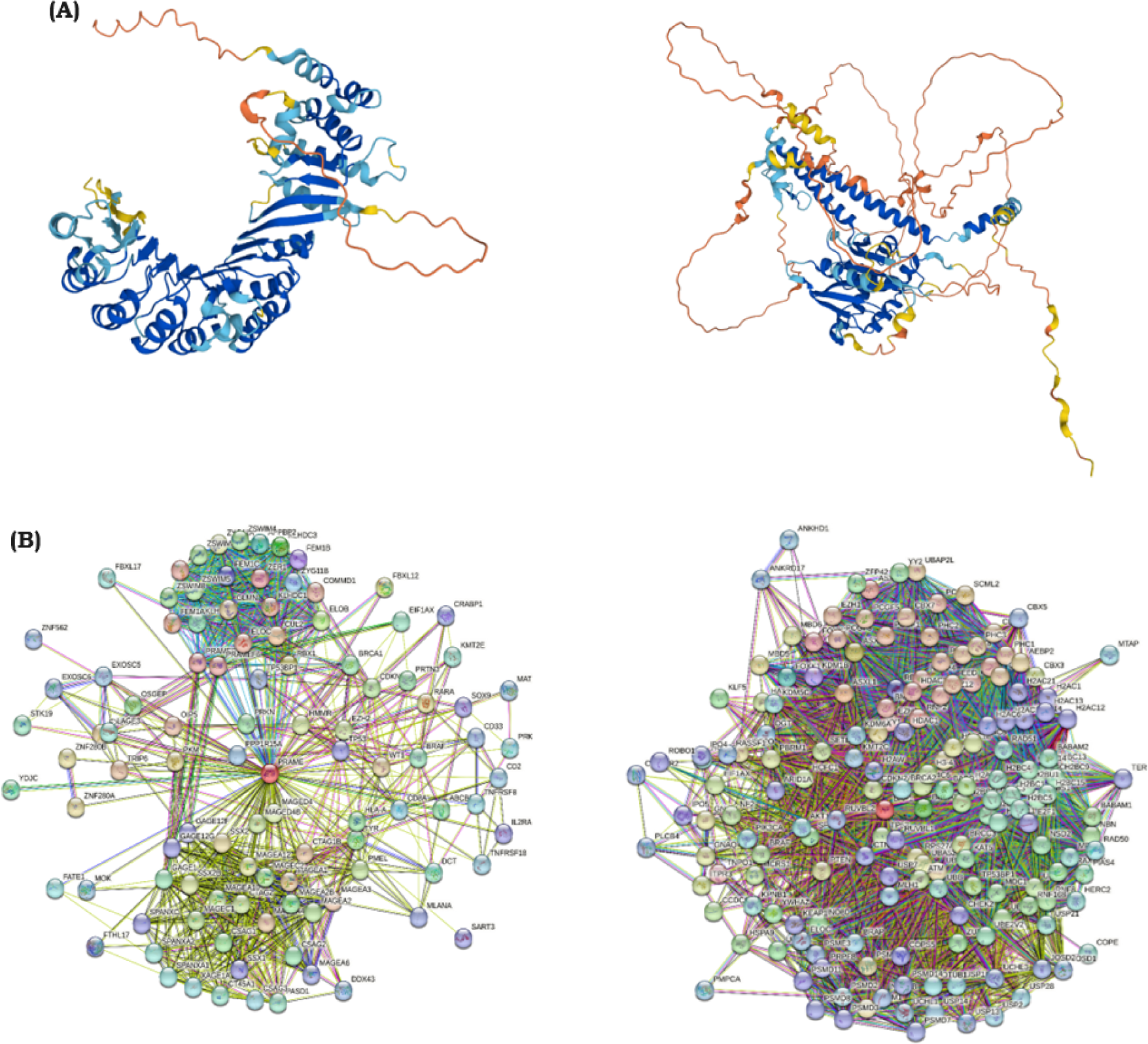
Evaluation of functional intrinsic disorder in human PRAME (UniProt ID: P78395) and human BAP1 (UniProt ID: Q92560). A. 3D structure of a protein modeled by AlphaFold2 This 3-D structure is color-coded based on the “Predicted Local Distance Difference Test” (pLDDT) values, where segments predicted with very high (pLDDT > 90), high (90 > pLDDT > 70), low (70 > pLDDT > 50), and very low (pLDDT < 50) confidence are shown by blue, cyan, yellow, and orange colors respectively [51]. B. STRING-generated protein-protein interaction network [52]. In both A and B, figures at left and right correspond to human PRAME and BAP1, respectively.

It is likely that these disordered/flexible regions of both proteins may play important functional roles. In fact, Figures 22B show that both PRAME and BAP1 contain multiple sites of different posttranslational modifications (PTMs), with BAP1 being heavily modified, especially its disordered regions. Furthermore, Figure 22B (bottom) shows that BAP1 is predicted to have 9 disorder-based binding sites known as molecular recognition features (MoRFs, residues 258-265, 289-306, 309-326, 330-401, 410-479, 497-508, 526-550, 555-575, and 673-684), which are disordered sub-regions that are not folded in the unbound state but may become ordered at interaction with specific partners. Figure 23B (right) represents STRING-generated protein-protein interaction (PPI) network entered at the human BAP1 and shows that this protein serves as a hub interacting with 173 proteins [47]. Overall, members of the resulting PPI network are highly interconnected, participating in 4526 interactions, and the average node degree of this network is 52. Despite its lower disorder content, human PRAME is also characterized by high binding promiscuity forming a PPI network containing 103 proteins connected via 866 interactions (Figure 23B (left)).

## 5. Discussion

Melanoma is one of the most prevalent skin cancers and affects nearly 325000 (as of 2020) people around the world every year [53]. PRAME, which is generally not expressed in tissues, is a tumor-associated antigen and shows expression specifically in cutaneous melanoma [54]. BAP1 is a tumor suppressor gene and plays an important role in regulation of the gene environment and its interactions.

In both PRAME and BAP1, there is a high degree of conservation of the residues, which can play an important role in protein function and interaction. In this study, we also found that there is sequence invariance, which can be indicative of potential function and evolutionary conservation. Previous studies have also shown that PRAME proteins, consists of Leucine-rich repeat (LRR) domains, which are composed of Leucine and Isoleucine residues and they play an important role in Protein-Protein Interaction (PPI) in different molecular pathways [55]. In BAP1, previous studies have also shown that the catalytic residues remain conserved [56]. Here we found that for PRAME there is an increase towards moderately flexible residues. Studies have shown that in PRAME, the regions with functionality are located in the Intrinsically Disordered Protein Regions (IDPRs) and are flexible [9]. In BAP1, there was an increase in the disorder promoting residues and a decrease in the order promoting residues and previous studies have also shown that the increase in the intrinsic disorder region helps in the establishment of a complex PPI module [13]. We observed that apart from Leucine, Glutamic acid poly strings of lengths 2 and 3 were also present in all the PRAME sequences. Furthermore, Glutamic acid poly strings of varied lengths were present in BAP1 sequences. This provides an important insite on the role of glutamic acid residues and how it can modulate the protein interaction. Previous studies have shown that Glutamic acid residues play a very important role as mutations in certain residues can lead to cancer [57].

A higher percentage of PP residue change was observed in BAP1, which can be correlated with the higher percentage of polar residues across the BAP1 sequences. Any changes in the distribution of the polarity change can lead to protein misfolding and thereby a disease phenotype [58]. Further studies in this direction can provide valuable information on how the polarity of amino acid residues can affect the protein folding, stabilization and interaction with other residues and proteins. It was found that for both PRAME and BAP1 sequences, there is a balanced distribution between the acidic and basic residues and any changes in the acidic and basic residue distribution can lead to changes in native structure, functional characteristics and interactions. Studies on acidic, basic, and neutral distributions are very essential as changes in the ratio of AB and BA can affect the interaction with other partners and can lead to gene dysregulation [59].

In this study we also found that for PRAME, glutamic acid had the highest median percentage of ‘D’ residues, indicating that Glutamic acid might be responsible for promoting disordered regions in PRAME sequences. For BAP1, Proline had the highest median percentage of ‘D’ residues and previous studies have shown that Proline content in BAP1 is more than the average proline content of the entire UniProt database [13].

Present study sheds light on crucial aspects of PRAME and BAP1 proteins, unraveling their structural and functional intricacies alongwith providing valuable insights into the potential roles of these proteins in melanoma and cancer-related processes. Furthermore, the findings contribute to our understanding of the potential impact on protein folding, stability, and interactions, emphasizing the importance of these factors in disease phenotypes. This investigation illuminates our comprehension of the molecular characteristics of PRAME and BAP1, providing a foundation for future research aimed at developing targeted therapies for melanoma.

## Acknowledgements

We all the authors would like to thank wholeheartedly Mr. Arindam Samanta for his insightful comments.

## Author contributions statement

SSH, DN, and AHJ conceived the problem and theoretical experiments. DN, SSH, VNU, AHJ, TB, and PB executed the results and performed the analyses. SSH, DN, VNU, AHJ, TB, EMR, and KL wrote the initial draft. All authors reviewed and edited the manuscript. All the authors checked, reviewed, and approved the final version of the manuscript.

